# The Mouse Inferior Colliculus Responds Preferentially to Non-Ultrasonic Vocalizations

**DOI:** 10.1101/2024.02.09.579664

**Authors:** Mahtab Tehrani, Sharad Shanbhag, Julia J. Huyck, Rahi Patel, Diana Kazimiersky, Jeffrey J. Wenstrup

## Abstract

The inferior colliculus (IC), the midbrain auditory integration center, analyzes information about social vocalizations and provides substrates for higher level processing of vocal signals. We used multi-channel recordings to characterize and localize responses to social vocalizations and synthetic stimuli within the IC of female and male mice, both urethane-anesthetized and unanesthetized. We compared responses to ultrasonic vocalizations (USVs) with other vocalizations in the mouse repertoire and related vocal responses to frequency tuning, IC subdivisions, and sex. Responses to lower frequency, broadband social vocalizations were widespread in IC, well represented throughout the tonotopic axis, across subdivisions, and in both sexes. Responses to USVs were much more limited. Although we observed some differences in tonal and vocal responses by sex and subdivision, representations of vocal responses by sex and subdivision were largely the same. For most units, responses to vocal signals occurred only when frequency response areas overlapped with spectra of the vocal signals. Since tuning to frequencies contained within the highest frequency USVs is limited (< 15% of IC units), responses to these vocalizations are correspondingly limited (< 5% of sound-responsive units). These results highlight a paradox of USV processing in some rodents: although USVs are the most abundant social vocalization, their representation and the representation of corresponding frequencies is less than lower frequency social vocalizations. We interpret this paradox in light of observations suggesting that USVs with lower frequency elements (<50 kHz) are associated with increased emotional intensity and engage a larger population of neurons in the mouse auditory system.

**SIGNIFICANCE STATEMENT:** The inferior colliculus (IC) integrates multiple inputs to analyze information about social vocalizations. In mice, we show that the most common type of social vocalization, the ultrasonic vocalization or USV, was poorly represented in IC compared to lower frequency vocalizations. For most neurons, responses to vocal signals occurred only when frequency response areas overlapped with vocalization spectra. These results highlight a paradox of USV processing in some rodent auditory systems: although USVs are the most abundant social vocalization, their representation and representation of corresponding frequencies is less than lower frequency social vocalizations. These results suggest that USVs with lower frequency elements (<50 kHz)—associated with increased emotional intensity—will engage a larger population of neurons in the mouse auditory system.

## INTRODUCTION

Vocal communication serves critical functions in the social interactions of many vertebrates. Vocal signals convey information about reproductive state, territorial dominance, aggression, affiliation, alarm, isolation, and complex cognitive processes (Gill and Bierema, 2013; Witzany, 2014). The brain’s analysis and representation of these signals derives from an array of processing steps beginning with the ear and continuing through and beyond ascending auditory pathways. Ultimately, this processing results in both internal and overt responses to the received vocal signals.

This study focuses on the representation of vocal communication signals in the inferior colliculus (IC), an auditory center within which many ascending, descending, and modulatory inputs converge and many complex tuning properties emerge (Winer and Schreiner, 2005; Wenstrup et al., 2012; Ito and Malmierca, 2018; Schofield and Hurley, 2018). As the nearly exclusive source of auditory projections to higher centers, the IC neural population response to vocal signals (Woolley and Portfors, 2013; Lyzwa et al., 2016) serves as the raw material for the processing in auditory thalamus and cortex, as well as extra-auditory regions such as the amygdala and striatum. The novel aspects of this study relate to its focus on broad features of responsiveness to vocal signals. How does the IC neuronal population respond to the diverse elements of an animal’s vocal repertoire? How does the population response relate to fundamental aspects of IC organization and function, such as subdivisions, tonotopic organization, and frequency tuning? How might vocal responses differ between females and males given some clear sex-based differences in vocal communication?

We consider these issues within the context of the mouse auditory system, a well-studied and complex communication system that also offers greater accessibility of experimental genetic tools (Brudzynski, 2018). In mice, ultrasonic vocalizations (USVs) are by far the most extensively studied type of vocal communication signal, due to their frequent occurrence and significance in courtship and other social behaviors (Sewell, 1972; Ehret, 2005; Holy and Guo, 2005; Portfors and Perkel, 2014; Sangiamo et al., 2020). One paradoxical aspect of USV processing in mice is that USV signals, with energy predominantly above ∼45 kHz, are processed by an auditory system with responsiveness primarily below 45 kHz (Egorova et al., 2001; Taberner and Liberman, 2005; Portfors, 2018). Several authors have suggested how this paradox may be resolved, with two broad hypotheses: 1) responses to USVs are carried by neurons tuned to lower frequencies (Portfors and Roberts, 2014; Portfors, 2018), or 2) there are in fact large numbers of neurons tuned to ultrasonic frequencies in the mouse IC (Garcia-Lazaro et al., 2015). We conducted this study to bring a population analysis to bear on the issue.

While the significance of USVs is clear, mice emit other vocal communication signals that have been less frequently studied, both in terms of their behavioral significance and their neural representations. These categories of mouse vocalizations, with lower fundamental frequencies and broader bandwidth, include the Low Frequency Harmonic call (LFH, i.e., the mouse squeak, 5-100 kHz), Mid-Frequency Vocalizations (MFV, 9-40 kHz), and Noisy calls (10-120 kHz) (Sewell, 1972; Portfors, 2007; Grimsley et al., 2011; Grimsley et al., 2016). They are emitted in several behavioral contexts such as mating, isolation, agitation, fighting or restraint (Grimsley et al., 2013; Grimsley et al., 2016; Keesom and Hurley, 2016), and evoke behavioral and physiological responses in listening mice (Niemczura et al., 2020; Ronald et al., 2020; Dornellas et al., 2021; Ghasemahmad et al., 2022). How are these vocalizations represented in the IC, and how do these representations compare to USVs?

To address these questions, we examined IC responses to a broad range of mouse vocalizations in both female and male mice. We specifically examined the relationship between vocal responsiveness of neurons and their frequency tuning properties. We further examined how responsiveness is distributed across the major IC subdivisions– central, external and dorsal–since these subdivisions project in partially distinct pathways through auditory thalamus to the cortex and amygdala (Wenstrup, 2005; Mellott et al., 2014). We predicted that population responsiveness to mouse social vocalizations differs among the subdivisions. Further, we predicted that responses to some vocal signals, especially USVs emitted during mating, differ between the sexes.

## MATERIALS AND METHODS

This study analyzed neuronal responses throughout the inferior colliculus (IC) to several categories of social vocalizations and other acoustic stimuli. Adult CBA/CaJ mice (Jackson Laboratory, Bar Harbor, Maine, USA) between the ages of 102 and 244 days were tested under urethane anesthesia (female: n=12, male: n=11) or while unanesthetized (females: n=3, males: n=6).

Mice were housed in our animal facility on a reversed, 12-hr light/dark cycle. Food and water were provided *ad libitum*. Animals were group housed in same-sex cages until the day of first surgery, after which they were singly housed. All procedures were approved by the Institutional Animal Care and Use Committee at the Northeast Ohio Medical University.

### Surgical and other Animal Procedures

At least two days prior to neurophysiological recordings, mice underwent surgery to cement a metal pin (headpost) onto the skull to facilitate restraint of the experimental mice during recordings. Mice were anesthetized with a mixture of isoflurane and O_2_ (2-2.5% flow rate). Ketoprofen (5 mg/kg, Sigma-Aldrich, Saint Louis, MO, USA) was subcutaneously injected to relieve pain and limit inflammation. Fur overlying the dorsal surface of the skull was removed using depilatory lotion. The skin was then disinfected with betadine and ethanol. An incision was made along the midline and the skin and muscle were retracted to expose the underlying skull. Next, the headpost was glued onto the skull using Metabond (Parkell Inc., Edgewood, NY, USA). A small craniotomy was made near the headpost through which a fine silver wire (reference electrode) was inserted into the cerebral cortex and fixed onto the skull using dental cement (sds Kerr, Orange, CA, USA). After surgery, local anesthetic (lidocaine) and antibiotic cream were applied to the surgical area and the mouse was returned to its home cage for recovery.

On recording days, a craniotomy (∼1.5 mm diameter) was made over the IC under isoflurane anesthesia and then covered with a silicone sealant (Kwik-Sil, WPI, Sarasota, FL, USA) until recording time. After mice were allowed to recover from surgery (∼30 min), most received urethane (total: 1 g/kg, Sigma-Aldrich, Saint Louis, MO, USA) administered in three intraperitoneal injections over a period of 1.5 hr. Depth of anesthesia was assessed by observing the animal’s reaction to experimenter handling. Once the anesthetic took effect (within a few minutes of administering the final dose), the mice showed reduced body and whisker movement and minimal to no startle. Nine animals received no urethane during recording sessions.

To determine the estrous stage of female mice, a sample of vaginal cells was obtained immediately following the last injection of urethane. Vaginal smear samples were collected using a small plastic transfer pipette filled with ∼100 µl of double-distilled water. The vaginal smear was then pipetted onto a clean, glass microscope slide and let dry on a slide warmer. Next, the specimen was covered with a few drops of Cresyl Violet Acetate staining solution, set aside for 10 min, then rinsed with double-distilled water and coverslipped. The estrous stage was identified microscopically based on the most abundant cell type in the sample: squamous epithelial cells (estrus), nucleated cornified cells (proestrus), or leukocytes (diestrus) as described by (McLean et al., 2012). The estrous stage was determined on each recording day. Neural recordings in the proestrous and diestrus stages were combined into a “non-estrus” category for response analyses.

### Acoustic Stimuli

Acoustic stimuli included noise and tone bursts as well as pre-recorded mouse vocal stimuli. Broadband noise (BBN) bursts had 4-80 kHz bandwidth, 100 ms duration, with 5 ms rise-fall times. Tone bursts ranged from 4-90 kHz, with 100 ms duration and 5 ms rise-fall times. BBN and tonal stimuli were repeated at 5/s. Pre-recorded stimuli (WAV stimuli) included 10 syllables representing distinct acoustic and behavioral categories of mouse vocalizations (Table 1). The syllables were chosen to represent a range of complexity and frequency among ultrasonic vocalizations (USVs) as well as lower frequency syllables including the LFH call, the noisy call, and mid-frequency vocalizations (MFVs) (Grimsley et al., 2011, 2016). We used multiple versions of the MFV call since its neural responses have not yet been investigated. The vocalizations were previously recorded from male-female pairs during mating interactions and males or females placed in a restraining jacket (Ghasemahmad et al., 2022). The vocal signals were recorded using condenser microphones (CM16/CMPA, Avisoft Bioacoustics, Berlin, Germany) connected to a multichannel amplifier and A/D converter (UltraSoundGate 416H, Avisoft Bioacoustics). Acoustic signals were digitized at 500 kHz, then conditioned (normalized, noise reduced, filtered) in Adobe Audition (2019). During playback, each vocal stimulus was preceded (45 ms, 5 ms rise time) and followed (30 ms, 5 ms fall time) by the background noise level present in the acoustic recording of the vocalization. The recording procedures and classification of the vocalizations are described in previous publications (Grimsley et al., 2016, Ghasemahmad et al., 2022). WAV stimuli also included BBN and tone bursts (70 and 80 kHz) for comparison to vocal stimuli. WAV stimuli were presented at 2/s.

**Table 1.**
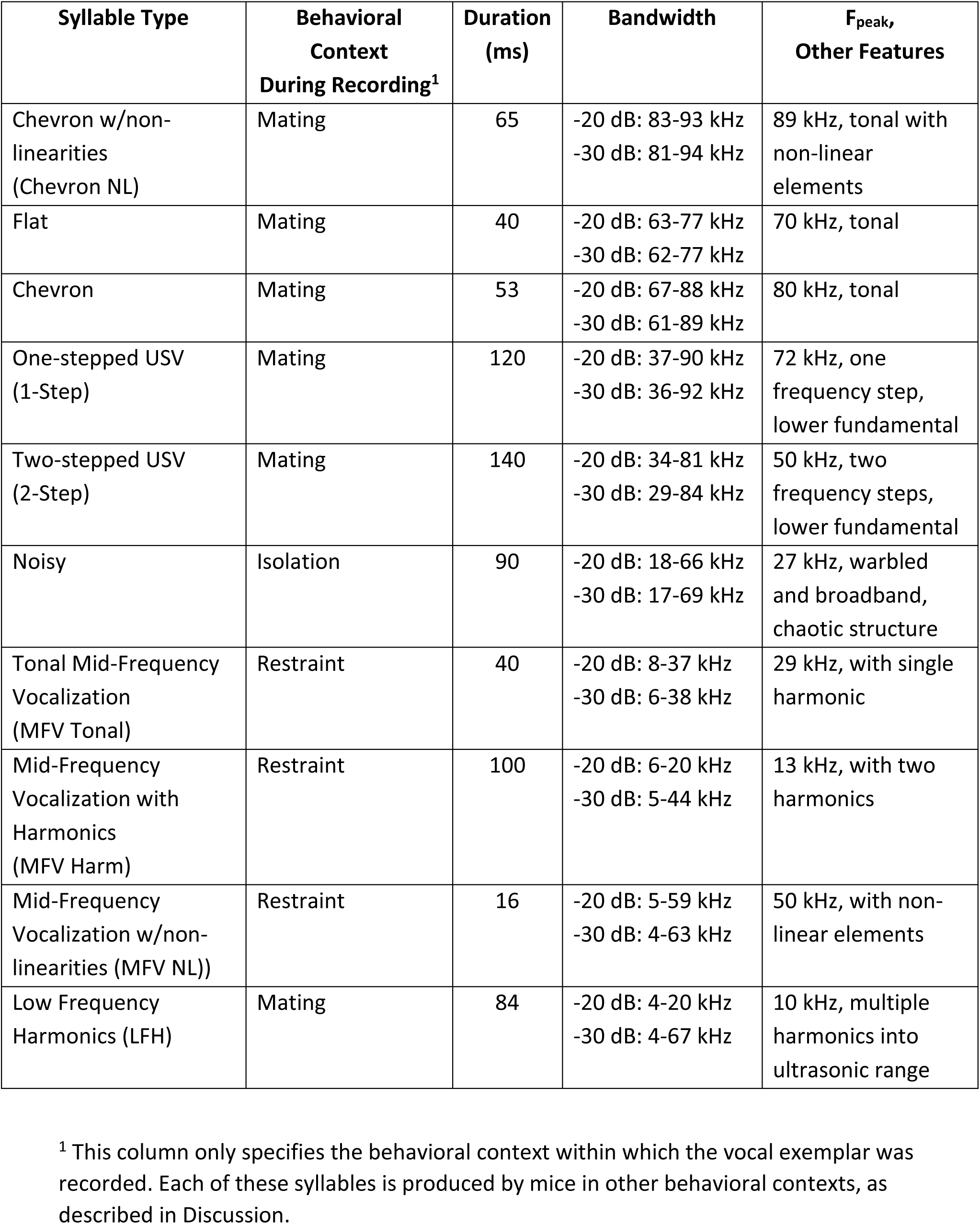
Acoustic features and associated behavioral context of vocal stimuli.

Digitized acoustic stimuli were converted to analog signals at 195.3 kHz and 16-bit depth using a TDT RZ6 system (Tucker-Davis Technologies, Alachua, FL, USA). Signals were amplified (Parasound Model HCA-750A) and sent to a ribbon tweeter loudspeaker (LCY-K100, Ying Tai Trading, Hong Kong). The frequency response of the audio D/A system was measured using a calibration microphone (Brüel and Kjaer 4939 ¼” microphone and preamplifier, Nexxus amplifier, Hottinger Brüel and Kjaer, Duluth, GA, USA) and stimuli were frequency compensated using custom Matlab software (Shanbhag, https://github.com/TytoLogy/NICal).

### Neurophysiological Recordings

Recordings were obtained from separate groups of urethane-anesthetized (ANEST) and unanesthetized (UNANEST) mice, placed inside a sound-attenuating chamber (Industrial Acoustics, Bronx, NY, USA). On the day of the experiment, the mouse was placed inside a plastic tube and head-fixed within a custom stereotaxic apparatus, then the silicon seal covering the IC and the dura were removed. A 4 X 4 multi-site, multi-shank neural probe (NeuroNexus A4×4-3mm-100-125-177-A16, Ann Arbor, MI, USA) pre-coated with DiI (Biotium, Fremont, CA, USA) for fluorescent detection of recording sites was advanced dorsoventrally at 5 µm/s through the IC using a hydraulic micropositioner (Model 650, Kopf Instruments, Tujunga, CA, USA). Extracellular signals were amplified and sampled at 48.8 kHz with 16-bit resolution (Model PZ2-32, RZ5D, Tucker-Davis Technologies, Alachua, FL, USA). Acoustic stimulation and data acquisition were controlled using custom MATLAB software. Sound was presented from a speaker placed 10 cm from the mouse’s head at a 45° angle into the sound field contralateral to the recording site. BBN (65-70 dB SPL) was used as a search stimulus as the recording probe advanced.

The goal of the recordings in ANEST mice was to map the distribution of responses with complete data sets in one or more penetrations per day. Such datasets were more difficult to obtain from UNANEST recordings. We aimed to sample responses throughout all dimensions of the IC, across the entire tonotopic range and including all major subdivisions. In ANEST recordings, the probe was initially advanced until auditory-evoked activity was encountered. Tests were then performed as described below. After auditory responses were thoroughly characterized at one probe depth, the probe was advanced ventrally by 400 µm for a new set of recordings. Overall, there were 2-5 steps of 400 um until the entire dorsoventral depth of the IC was characterized. This was verified by ensuring that the most ventral sites at the greatest depth displayed no auditory responses. After each advance of the probe, we allowed ∼10 min for neural activity to stabilize prior to the start of recording.

The goal of UNANEST recordings was to examine whether frequency tuning properties or vocalization responses were affected by anesthesia. Since our interest here was primarily in the responses of neurons tuned to lower frequencies (<30 kHz), we used a slightly different approach to sampling neural activity. Here, the probe was advanced until strong auditory-evoked activity occurred on all channels. These were typically concentrated in the dorsal half of IC and there was no attempt to systematically sample throughout IC. Testing was identical to that performed during ANEST recordings (see below), except that the data were obtained at only one probe depth. In all experiments, the animals were sacrificed after the final recording session and brains were processed for histological analysis (see below).

The following tests were performed at each recording location: 1) rate-level function in response to BBN (0-70 dB SPL in 10 dB steps, 20 repetitions); 2) Frequency Response Area (FRA) (4-90 kHz in 1/3 octave steps, 10-80 dB SPL in 10 dB steps, 20 repetitions); 3) responses to the 13 WAV stimuli at peak levels of 20, 40, 60 and 70 dB SPL. At each sound level, WAV stimuli were repeated 20 times in a blocked-randomized order where a block consisted of a random sequence of stimuli with no repeats presented once, followed by another block, and so on. A “null” or no-sound interval was also included to assess background firing.

### Spike Data Analyses

Plexon Offline Sorter version 3 (Plexon, Dallas, TX, USA) was used to sort the recorded neural data. Raw data were bandpass filtered (300 – 4,000 Hz) prior to spike sorting. Using spike features and principal component analyses, spikes were clustered as multi-units.

#### Spike Density Functions

For each unit, spike trains in response to each stimulus were converted to a mean spike density function (SDF) with 1 ms time bins. Single-trial spike trains were first convolved with a Gaussian kernel with 10 ms width. The MATLAB bootci function was then used to compute a bootstrap estimate of the mean SDF and 95% confidence intervals (CI). Background (BG) firing rates and the 5^th^ and 95^th^ percentiles of the BG firing distribution were estimated from this bootstrap analysis of the “null stimulus” SDFs.

#### Unit responses

Unit responsiveness to vocal stimuli was determined by comparing the stimulus-response SDF with the BG firing. For each stimulus, an analysis window was defined as the time from stimulus onset to 45 ms after stimulus offset. This post-stimulus time was selected to account for any firing changes after stimulus offset. Significant increases in firing were classified as points where the lower 95% CI of the stimulus-response SDF was greater than the 95^th^ percentile of the BG firing distribution. Similarly, significant decreases in firing were points where the upper 95% CI of the stimulus-response SDF was lower than the 5^th^ percentile of the BG firing distribution. Units were classified as being responsive to a given stimulus if there was a significant increase or decrease in firing for a minimum duration of 10 ms within the vocal stimulus window. Stimulus windows were defined as the time between stimulus onset and offset (see Table 1 for stimulus durations for vocal stimuli).

#### Determination of FRA and CF

For each combination of tone frequency and level, the mean number of spikes within the 100 ms stimulus duration was calculated. A custom MATLAB program then displayed each unit’s FRA as a pseudocolor plot and surface. From this display, a user selected the point on the FRA plot that corresponded with the lowest level that elicited firing at a specific frequency. Units that showed no response to tones or no frequency tuning were classified as having no CF.

#### Calculation of frequency tuning bandwidth

A bootstrap procedure was used to determine the frequency tuning bandwidth (BW) of units from FRA data at 60 and 70 dB. For units with a defined CF, a random sample of 20 spike counts was drawn (with replacement) at each tested frequency at the desired stimulus amplitude. Using the MATLAB makima function, an interpolated curve with 100 points was fit to these data. This procedure was repeated 1000 times and a bootstrap mean frequency tuning curve (FTC) and 95% CI were computed from these bootstrapped fits. The background rate of the unit was determined from the 0 dB stimulus data and a bootstrapped mean and CI was then determined using the MATLAB bootci function. The lower frequency limit of BW was then set to the lowest frequency at which the lower CI of the FTC was greater than the upper CI of the BG mean. Similarly, the upper frequency limit of BW was set to the highest frequency at which the lower CI of the FTC was above the upper CI of the BG firing.

#### Calculation of overlap between unit tuning and call spectrum

Unit frequency tuning bandwidth and the spectral energy of the stimulus were used to calculate the bandwidth-spectrum overlap of each unit to each stimulus (Fig. 8A). Overlap was defined as the difference between the upper (high frequency) limit of frequency tuning of a unit and the lower frequency limit (at -30 dB) of the spectrum of a call. Positive overlap values therefore indicate the amount of congruence between the unit bandwidth and the stimulus spectral energy, while negative values indicate the distance between the upper tuning limits of the unit and the lower frequency limit of the stimulus.

### Statistical Analyses

#### CF distributions

CF distributions were compared across males and females within the entire IC (Fig. 2A) and in each subdivision (Fig. 2B-D) using a Chi-square goodness of fit test (MATLAB chi2gof function).

#### Bandwidth distributions

The distributions of the upper frequency limits for IC units were analyzed across CF groups and IC subdivisions (Fig. 3B-D), and within CIC across sex and estrous stage (Fig. 3E) using Independent-Samples Kruskal-Wallis tests (Tables 3-1, 3-2). These analyses were performed using SPSS (version 28, IBM Corp., 2021). All *post-hoc* tests used Benjamini-Hochberg corrections with a False Discovery Rate of 0.05. This correction method assigns a different critical p-value for each of the comparisons, based on the rank of each p-value when the p-values for all comparisons are ordered from smallest to largest. If the raw p-value is less than the critical p-value for that comparison, then the result of the analysis is significant (Benjamini and Hochberg, 2000).

#### Vocal response distributions

Responsiveness of IC subdivisions to the vocal categories (Fig. 10) was analyzed using a Linear Mixed Effects Model (LMEM) using the lme4 package (Bates et al., 2015) in R (R Core Team, 2021; https://www.R-project.org) in an R Studio environment (R Studio, n.d.; http://www.posit.co). Data were analyzed using a full-factorial mixed effects model with cell count as the dependent variable. The fixed effects were IC region (nominal) and vocalization category (nominal) and the interaction between the two. To account for between-unit variability, the intercepts were allowed to vary randomly across unit ID numbers (N = 1149). A Poisson distribution was used in the LMEM because the data were count data. All *post-hoc* tests used Benjamini-Hochberg corrections with a False Discovery Rate of 0.05. A similar analysis revealed no sex differences in responses of CIC units to the vocal categories.

### Histological Verification of Recording Locations

The DiI-illuminated probe tracks were photographed using a SPOT RT3 camera and SPOT Advanced Plus imaging software (version 4.7) mounted on a Zeiss Axio Imager M2 fluorescence microscope (Fig. 1A, B). The recording sites were histologically reconstructed first by identifying the brain section containing the most intense DiI labeling as the penetration site. The rostrocaudal (R-C) recording location was determined by first identifying the sections containing the most rostral and most caudal extents of the IC using Nissl-stained images and the mouse brain atlas (Franklin and Paxinos, 2008). Each section through the IC including that containing the recording location was then assigned a percentage across the R-C dimension of the IC. The mediolateral (M-L) location of each shank was determined by measuring the distance from the midline to either the most medial or lateral shank track using the image analysis software. The M-L positions for the other three shanks were calculated based on the known inter-shank distance of 125 µm. The M-L measurements were then superimposed onto the atlas section corresponding to the fluorescent image containing the probe tracks. Finally, the dorsoventral (D-V) position for each recording site and thus the IC subdivision assignment was determined based on the known recording probe inter-site distance of 100 µm and the overall depth of recording. A small number of units were identified as “border” units, being located on the borders between subdivisions. These were excluded from subdivision-based analyses of response properties.

**Figure 1.**
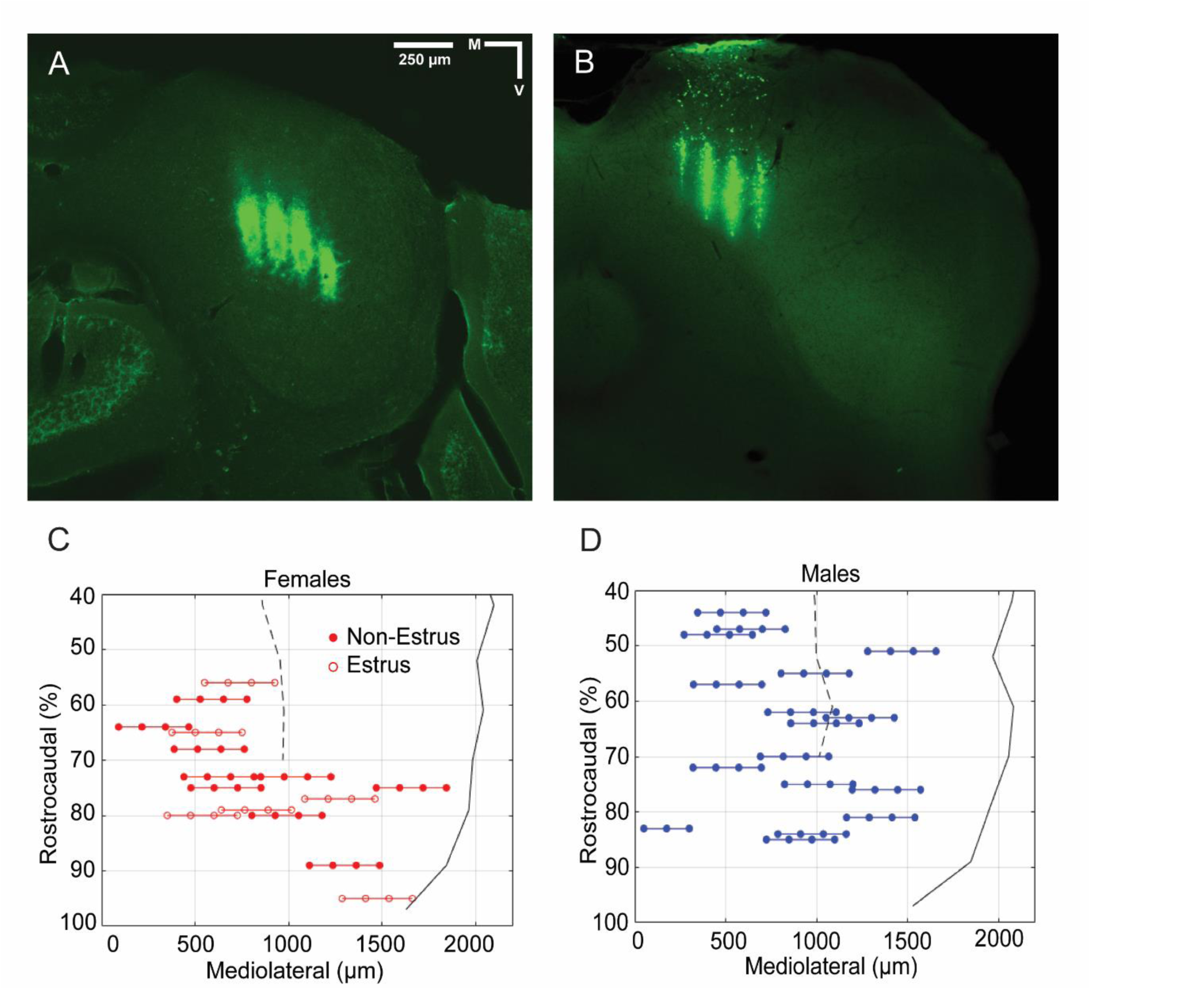
Histological reconstruction of IC recording sites. **A, B.** Photomicrographs show Di-I labeling from multi-shank electrodes through the central (CIC) and dorsal cortex (DCIC) of the IC, respectively. **C, D.** Reconstructions of all recording locations in females and males, respectively. Colored lines connect the four shanks of recording electrodes which are represented by filled (non-estrus females and males) or open (estrus females) circles. Dashed black lines show the lateral border of the periaqueductal gray (PAG), while solid black lines show the lateral border of the IC. There is extensive overlap of recording sites in DCIC and CIC of female and male subjects, but less overlap in external cortex of the IC (ECIC) recordings. The rostral 40% of IC contains ECIC but no DCIC or CIC; it is inaccessible in dorso-ventral penetrations due to a sinus overlying the rostral IC and superior colliculus.

The distribution of histologically reconstructed recording sites is shown in Figure 1C, D. Electrodes were placed to record from the rostro-caudal and medio-lateral extents of the mouse IC. Due to the presence of a sinus over the rostral IC and superior colliculus, we were unable to place electrodes in the rostral 40% of IC. This part of IC includes the ECIC and little, if any, of CIC or DCIC.

## RESULTS

This study examined the sound-evoked responses of multi-unit recording sites (“units”) distributed within the IC of 12 female and 11 male ANEST mice (Fig. 1). Of 1,389 units localized to the IC, 1,212 were responsive to tones, noise, or vocal stimuli in our tests. We first describe frequency tuning among units in the IC sample, then responses across four categories of social vocalizations and the relationship between frequency tuning and vocal responses. Throughout, the auditory responses were analyzed by IC subdivision, by sex, and by female estrous state. To examine potential effects of anesthesia on vocal responses, additional recordings were obtained from UNANEST mice (3 females and 6 males). These data are described at the end of the Results and in Figures 10-12.

### Frequency Tuning

Among IC units, 1163 displayed organized frequency response areas (FRAs) with defined characteristic frequencies (CFs). Of these, 535 units were from females and 628 units were from males. The peak in the distribution of CF values across the sample was in the 13-16 kHz range, with a secondary peak near 51 kHz (Fig. 2A). Less than 10 units had CFs above 60 kHz. Based on a Chi-square analysis, the female and male distributions were significantly different (*X^2^* (9,628)=85.57, p< 0.001). Figure 2A shows that the peak CF was lower in frequency for females (13 kHz vs. 16 kHz in males), that the secondary peak at 51 kHz was substantially larger in males, and that only males had CFs of 64 kHz.

**Figure 2.**
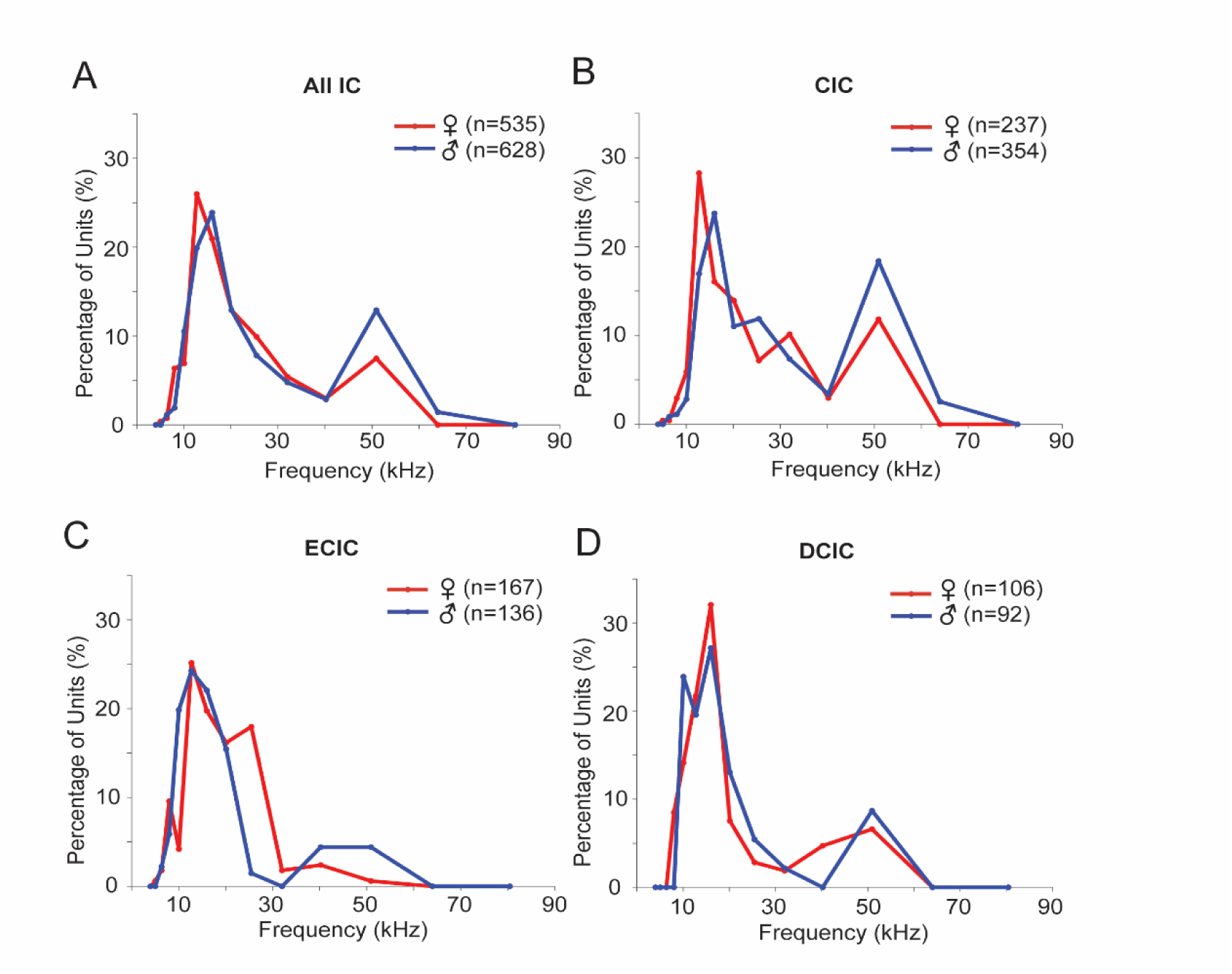
Characteristic frequency (CF) distributions across IC unit populations. **A.** CF distribution across entire IC sample shows the primary peak at 13-16 kHz and a secondary peak near 51 kHz. Female and male distributions are significantly different, particularly in the size of the secondary peak: *X^2^* (9,628)=85.57, p< 0.001. This sample includes all units for which a CF could be identified, including those for which the subdivision of IC is uncertain. **B.** CF distribution across the CIC sample in females and males. Similar to **A**, the female and male distributions significantly differ: *X^2^* (8,354)=78.60, p< 0.001. **C.** CF distribution across the ECIC sample in females and males. Compared to the overall IC and CIC distributions, there are fewer units tuned to frequencies above 32 kHz. These distributions differ in females and males: *X^2^* (6,136)=106.96, p<0.001. **D.** CF distribution across the DCIC sample in females and males. There is no significant difference in these distributions: *X^2^* (4,92)=4.63, p>0.05.

The CF distributions across subdivisions displayed noteworthy differences. The central IC (CIC) distributions displayed the greatest similarity to the overall IC distributions, which is not surprising since the CIC units formed the largest group (Fig. 2B). Moreover, the female and male distributions in CIC were significantly different (*X^2^* (8,354)=78.60, p< 0.001), with the same features as the overall distribution. That is, in males, the peak CF was higher than in females (16 kHz vs. 13 kHz), the secondary peak at 51 kHz was larger, and some units had CFs at 64 kHz (none in females). These 64 kHz sites in CIC were localized to ∼70% of the rostrocaudal extent of the IC and a depth of 1100-1500 μm.

CF distributions in the external IC (ECIC) differed from the CIC distributions, with substantially fewer units tuned to frequencies above 30 kHz (Fig. 2C). This was especially true for females, which displayed very few CFs above 30 kHz. The female and male distributions in ECIC differed significantly (*X^2^* (6,136)=106.96, p<0.001). In the dorsal cortex of IC (DCIC), the CF distributions were similar to the overall and CIC distribution, though with fewer high-CF units; a Chi-square analysis revealed no female/male difference (*X^2^* (4,92)=4.63, p>0.05).

We next examined the bandwidth of IC tonal responses to set the stage for later comparisons to units’ responses to vocal stimuli. Using 60 dB SPL tonal stimuli, we created a frequency response curve (FRC) from which we measured the lower and upper frequency limits. We then plotted the resulting bandwidths as a function of unit CF, and, within CF, by the upper limit of the FRC (Fig. 3A). This display, in which each horizontal line represents the range of frequencies to which one unit responds, reveals that most neurons responded to tonal stimuli above 8 kHz. Further, neurons tuned below 40 kHz displayed a very broad range of 60-dB SPL bandwidths. For example, among units with CFs of 13 kHz, lower-upper frequency limits of units ranged from 4-13 kHz to 4->64 kHz. Units with CFs above 32 kHz had broad bandwidths, nearly all responding to 8-51 kHz tones. Very few units responded to tones at 80 kHz or above.

**Figure 3.**
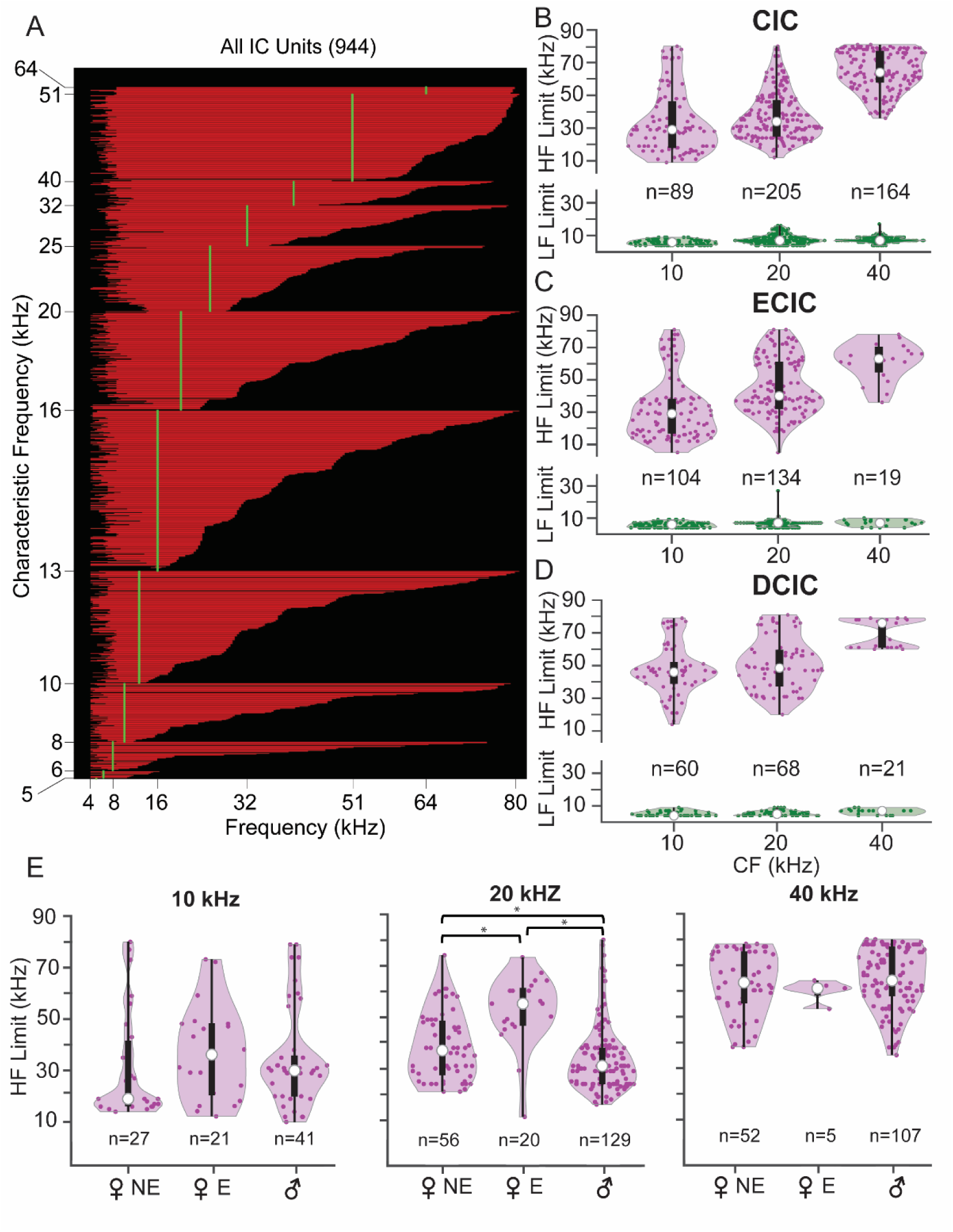
Bandwidth of frequency tuning varies substantially among IC units. **A.** Frequency response curves displayed as bandwidths (BW) obtained from responses of IC neurons to tones at 60 dB SPL. Unit BW is displayed as the extent of red line along x-axis (one unit per horizontal line). Units are sorted vertically by CF, then within CF by the upper frequency limit of the BW at 60 dB SPL. BWs varied widely among most CF groups, especially at lower CFs. **B-D.** The lower (green) and upper (purple) frequency limits of BWs displayed as violin plots in three octave-wide frequency bands, centered at 10, 20 and 40 kHz in CIC (**B)**, ECIC (**C**), and DCIC (**D**). **E**. Distribution of upper frequency limits across CF band, sex and estrous status within CIC. For all violin plots, open circle indicates median, bar indicates interquartile range. Asterisks show statistically significant comparisons (p<0.05). Detailed statistics in Tables 3-1, 3-2.

The lower and upper frequency limits of FRCs were then analyzed within three, octave-wide CF bands centered at 10, 20, and 40 kHz, and across IC subdivisions (Figs. 3B-D, Table 3-1). Across CF bands and subdivisions, there was no difference in the lower frequency limits (*in green*). For upper frequency limits (*in magenta*), we observed some differences within and across subdivisions. Within subdivisions, the expected trend was that higher CF units would display higher upper frequency limits. In agreement, 40 kHz CF bands always had the highest upper frequency limits (p<0.001). Unexpectedly, the upper frequency limits of units in the 10 kHz and 20 kHz bands were similar for both CIC and DCIC (p=0.07 and p=0.15, respectively). That is, units with CFs in these two bands were equally likely to respond to high frequency stimuli.

We next compared the upper frequency distributions across subdivisions (Table 3-1). In the 10 kHz band, DCIC units had significantly higher upper frequency limits (median 46 kHz) than ECIC or CIC units (median 29 kHz for each; p<0.001). At 20 kHz, all three subdivisions differed in their upper frequency distributions, with DCIC having the greatest (median 48.5 kHz; vs. CIC: p<0.001; vs. ECIC: p=0.03) and CIC the smallest values (median 34 kHz; vs. ECIC: p<0.001). This indicates broader frequency tuning in DCIC compared to the other subdivisions among units in the 10 and 20 kHz bands (Fig. 3D). At 40 kHz, the three subdivisions did not significantly differ in the distribution of the upper frequency limits of their tuning bandwidths (p=0.09).

Finally, we examined whether upper frequency limits of IC neurons showed sex- or estrous-related differences (Fig. 3E). We focused on the CIC, because only this subdivision had highly overlapping recording sites between the sexes and sufficiently large numbers of units from each sex. The main findings were that unit distributions of upper frequency limits in the 20 kHz CF band differed significantly by sex and estrous state. Median values in male units (31 kHz upper limit) and non-estrus female units (38 kHz) were much lower than in estrus female units (56 kHz) (p<0.001) (Table 3-2). Male and non-estrus female distributions at 20 kHz were also significantly different (p=0.03). For units in the 10 kHz and 40 kHz bands, there were no significant sex or estrous state differences (p=0.33 and p=0.23, respectively). These results suggest that, in the 20 kHz CF band, units in females (especially estrus females) were more likely to respond to higher frequency USVs than in males.

### Responses to Social Vocalizations: Relation to Frequency Tuning

We tested responses to 10 social vocalizations, including five USVs and five calls with fundamental frequencies at or below 20 kHz. In addition, we also tested responses to broadband noise (BBN) and tone bursts corresponding to peak frequencies within the flat (70 kHz) and chevron (80 kHz) USVs. Responses to these acoustic stimuli are displayed as both raster plots and Spike Density Functions (SDFs, Fig. 4). Units were considered to show a response to a stimulus by reference to the mean background discharge assessed in presentations of a “null” stimulus (see Materials and Methods).

**Figure 4.**
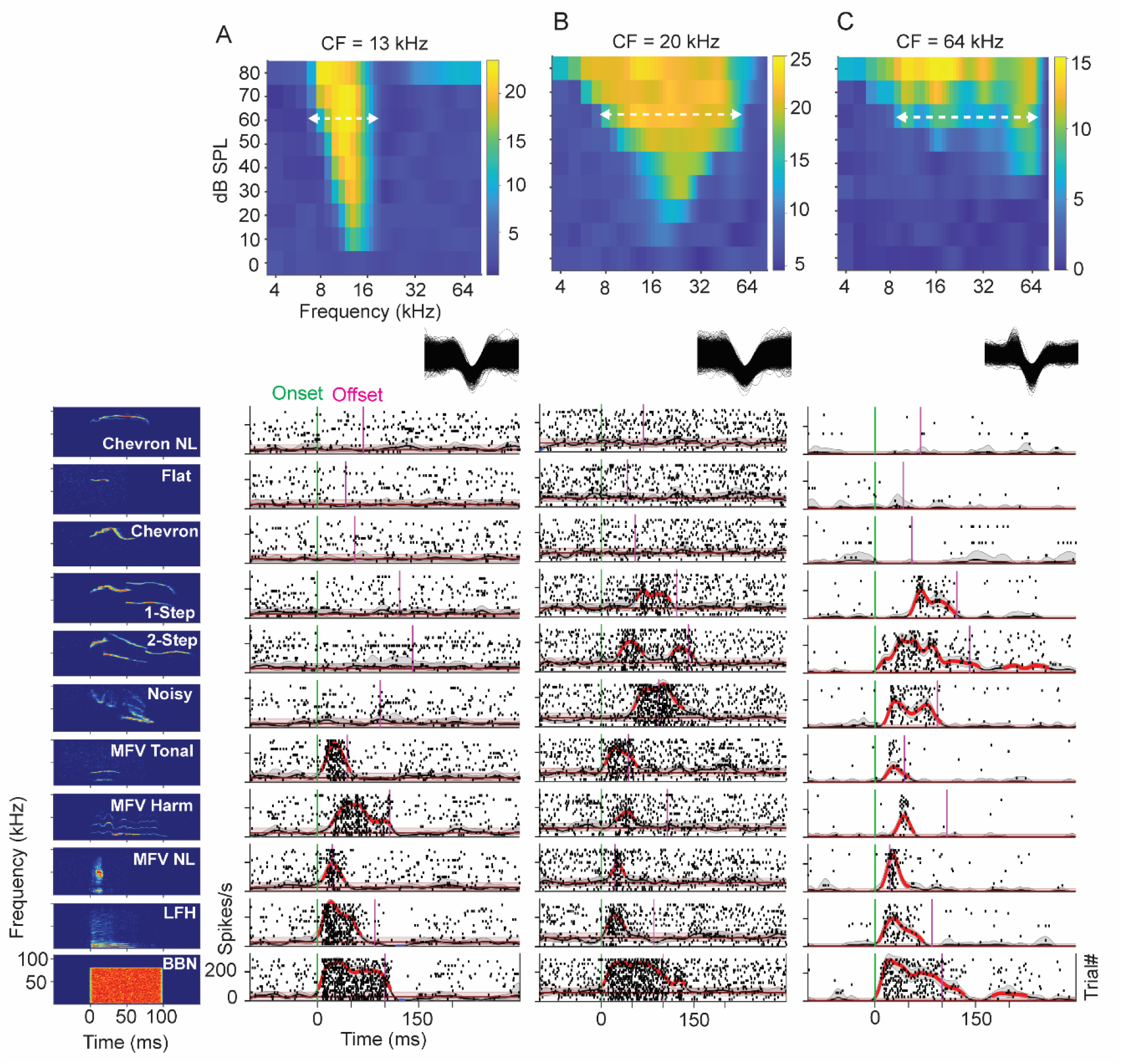
Example unit responses to each vocal stimulus: effect of frequency tuning and bandwidth. Spectrograms of BBN and 10 vocal stimuli (at left) are arranged vertically in order of increasing minimum frequency of stimulus. All stimuli presented at 60 dB SPL peak. Responses correspond to stimulus at left of row. **A**. Frequency response area (FRA) (top), overlaid spike waveforms (middle), and vocal responses (bottom) of unit with low CF (13 kHz) and 10 dB SPL threshold. Tuning width at 60 dB SPL was 7-19 kHz (horizontal white dashed line). Due to low CF and narrow frequency tuning at 60 dB SPL, responses of this unit are restricted to BBN, LFH and MFVs only. **B**. FRA of unit with somewhat higher CF (20 kHz) and threshold near 10 dB SPL, but greater bandwidth than unit A (7-62 kHz). Unit responds to stepped USVs but not to tonal USVs. **C**. FRA of unit with high CF (64 kHz) and threshold near 30 dB SPL (top) with vocalization responses (bottom). At 60 dB SPL, this unit has a broad tuning width (6-79 kHz, dashed white line). Due to high CF (64 kHz) and broad frequency response at 60 dB SPL, this unit responds to BBN and all categories of vocalizations except for tonal USVs (Chevron and Flat calls). Unit responses are displayed as spike rasters (black dots) and spike density functions (SDFs). Spike waveforms are 1 ms duration. SDF time bins significantly above mean background (BG) are highlighted in red. Confidence intervals are shown for SDF functions (grey shading) and for no-stimulus control (red shading).

Figure 4 shows the relationship between frequency tuning and responses to the vocal stimuli (at peak level of 60 dB SPL) in three units differing in CF and width of tuning. For a unit with CF of 13 kHz (Fig. 4A), threshold sensitivity was near 10 dB SPL and responsiveness at 60 dB SPL extended from 7-19 kHz (dashed line). Consistent with its FRA, the unit responded well to broadband noise and to vocal stimuli with energy below 16 kHz, but not to stimuli––either broad band or narrow band––with energy exclusively above 16 kHz. This includes stepped or frequency modulated signals that might be expected to generate lower frequency cochlear distortion products. For a unit with somewhat higher CF (20 kHz, threshold 10 dB SPL, Fig. 4B), the width of tuning at 60 dB SPL was much broader than in Figure 4A, extending from 7 to 62 kHz (dashed line). This unit responded to BBN and most vocalization types, likely the result of the unit’s broad tuning that encompasses frequencies within these vocalizations. However, the unit did not respond to the tonal USVs (chevron and flat calls), likely due to the high frequency content of these calls (≥ 70 kHz). For a unit with high-CF (64 kHz, threshold 30 dB SPL, Fig. 4C), the width of tuning at 60 dB SPL was similar to the 20 kHz unit in Figure 4B, from 6 to 79 kHz (dashed line). Like the unit in Figure 4B, this unit responded to BBN and most vocalization types, except the tonal USVs (chevron and flat calls). This was likely due to the high frequency content of these calls (≥ 70 kHz). While each of these calls contains some energy at the unit’s CF, the level was likely too low to evoke excitatory responses.

Features of IC vocal responses are revealed in population displays that plot the time course of excitatory and inhibitory responses to BBN and five of the vocal stimuli. In each plot (Fig. 5), responses of 944 IC units are arranged vertically according to ascending CF, then by increasing tuning bandwidth within each CF group, corresponding to Fig. 3A. Population responses for other stimuli are displayed in Fig. 5-1.

**Figure 5.**
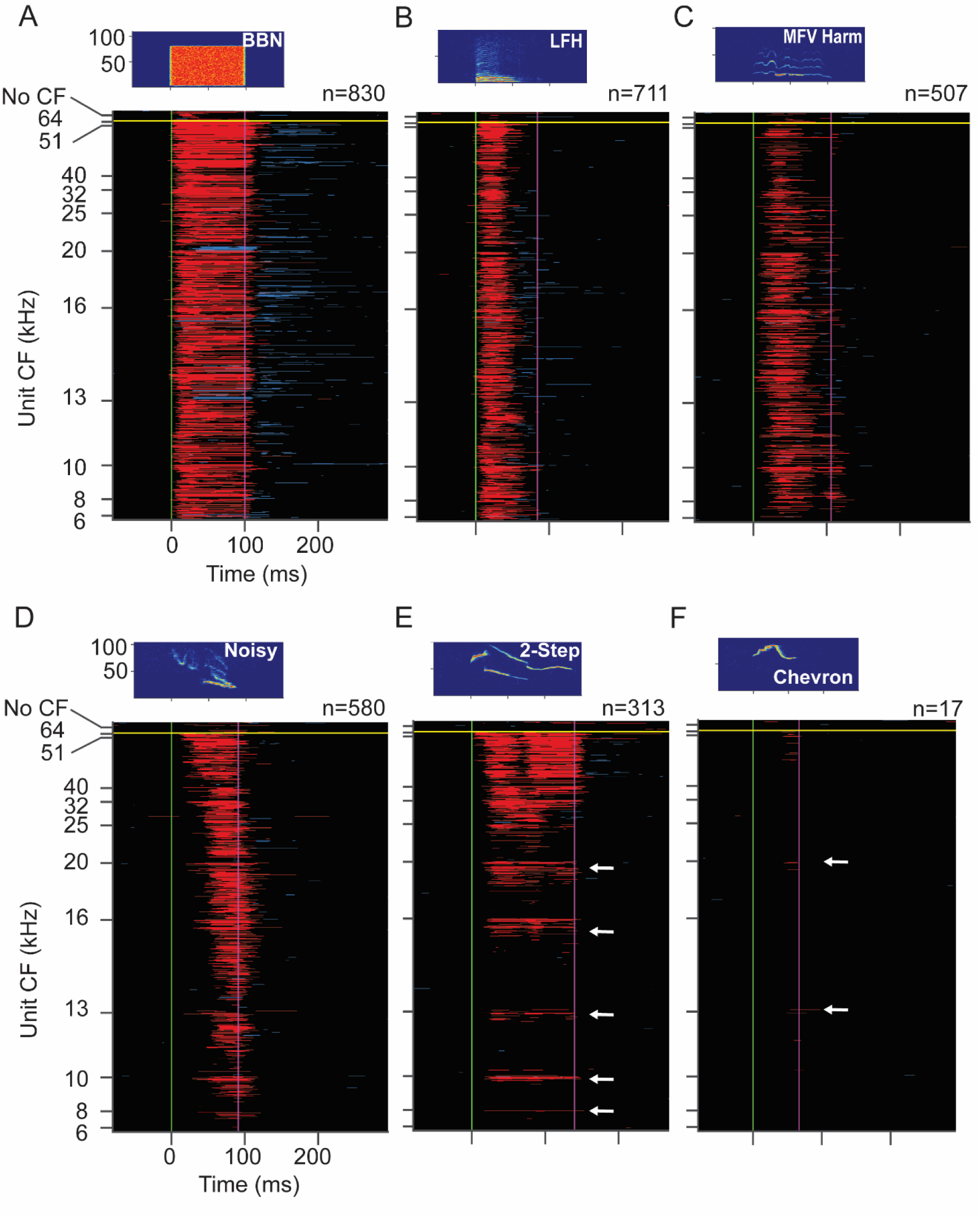
Responses of IC population to WAV stimuli as function of CF. **A-F.** Population response to the WAV stimulus indicated in the sonogram. Each plot shows excitatory (red) and inhibitory (blue) SDF responses to stimuli at 60 dB SPL peak. Each horizontal line represents response of one unit. Units are arranged vertically in order of increasing CF, and within each CF group in order of increasing upper limit of frequency tuning. Responses of units with no observable CF are placed above horizontal yellow line. Neon green and purple vertical lines mark onset and offset of each stimulus, respectively. Each plot shows the same 944 units, the subset of 1212 sound-responsive IC units that responded to at least one of the 60 dB SPL WAV stimuli. These units also illustrated in Fig. 3A. Stimulus spectrograms and number of responding units shown above main plot. **E-F.** White arrows indicate some lower CF units that respond to USVs. These are typically units with highest upper frequency limit in each CF group. Responses to other vocalizations in Fig. 5-1.

The BBN stimulus (Fig. 5A), as expected, evoked excitatory responses (in red) among most IC neurons with CFs from 5 to 64 kHz (830 of 1212 sound-responsive IC units, 68%). A limited number of units displayed predominantly inhibitory responses (in blue) during the stimulus. Excitatory responses were typically sustained, with many units showing long lasting inhibition after stimulus offset.

The LFH stimulus features low peak frequency (10 kHz) and -30 dB bandwidth of 4-67 kHz (Fig. 5B). The IC population response was somewhat less (59% of units) than to BBN, but still distributed across the entire range of unit CFs (Fig. 5B). The shorter duration responses to the LFH, compared to the BBN, may result from the shorter duration of the higher frequency harmonics in the LFH call. As with the response to BBN, several units showed inhibitory responses immediately after the excitatory responses ended.

The MFV Harmonic call has a higher fundamental frequency than BBN and the LFH call (Fig. 5C), with peak frequency of 13 kHz and -30 dB bandwidth of 5-44 kHz. It evoked responses in 42% of IC units from across the tonotopic range. Inhibitory responses sometimes followed the excitatory response, occurring well before stimulus offset.

Temporal patterns, especially response onsets, were variable, likely because peak energy in different frequency bands occurred at different times. Other variants of the MFV call evoked somewhat less (MFV NL, Fig. 5-1A) or somewhat more (MFV Tonal, Fig. 5-1B) responses among IC units.

The Noisy call has energy in a very broad band, from 17-69 kHz. Initially, call energy is in the ultrasonic range, while energy near 20 kHz occurs later in the signal. Two features of the IC population response are noteworthy. First, many units across the entire tonotopic range responded to this call (48%), including units with CFs below 16 kHz (Fig. 5D).

Although the Noisy stimulus contains no significant energy in this frequency band, a substantial number of units with CFs of 16 kHz or less responded. Second, the timing of responses shows a clear relationship to CF. High-CF units, responsive to high frequencies that occur early in the Noisy call, typically responded with the shortest latencies. Low-CF units responded with longer latency, most likely due to the later occurrence of lower frequencies in the Noisy stimulus. Inhibitory responses were fewer among this population.

USVs have energy entirely within the ultrasonic range but have different spectral features that generate very different population responses in the IC (Figs. 5E,F). The 2-stepped USV, with energy extending from 29-84 kHz, excited 26% of units, most tuned above 24 kHz. There was also a significant number of responses to this USV among neurons tuned below 24 kHz, clustered among units with higher upper frequency limits (Fig. 5E, arrows). Inhibitory responses were infrequent during the stimulus. Temporal patterns were frequency dependent. Only the highest CF units (CF = 64 kHz) displayed very short latency excitatory responses, likely initiated by the ∼62 kHz initial segment of the call. Higher CF units were also the only units that showed long lasting inhibition following the excitation. Excitatory responses of lower CF units often began following the initial step to the call’s lower frequency element near 42 kHz. The 1-stepped call, with somewhat higher minimum frequency, evoked a slightly lower population response, 19% of units (Fig. 5.1C).

Few IC units responded to tonal USVs or corresponding tone bursts. The chevron, with a bandwidth of 61-89 kHz, evoked responses in 1.4% of units when presented at the peak level of 60 dB SPL. Responsive units were mostly tuned above 50 kHz, but a few excitatory responses were seen among units with CFs below 25 kHz (Fig. 5F, arrows). Inhibitory responses were rare. Most responses began later, corresponding to the downward frequency modulation in the chevron signal. The flat syllable similarly evoked responses in very few units (1.1%), nearly all with CFs above 50 kHz (Fig. 5-1D). A 70 kHz tone burst, corresponding to the peak frequency of the flat call, also evoked minimal responses (Fig. 5-1E). The Chevron NL syllable, with the highest minimum and peak frequencies (81 and 89 kHz, respectively) and sideband non-linearities, evoked IC responses in two units at 60 dB SPL (Fig. 5-1F).

Figure 6A compares responses of the IC population to each of the WAV stimuli (at 60 dB SPL peak). Consistent with frequency tuning (Figs. 2, 3), the IC population responded well to broadband signals with lower frequency components, reasonably well to USVs with lower ultrasonic frequencies, but very poorly to USVs or tones with frequencies only > 60 kHz. This pattern of responsiveness, in which no vocal stimulus excites more than 60% of sound-responsive IC units, implies a degree of selectivity. Figure 6B displays selectivity, expressed as the number of vocal stimuli (maximum = 10) to which a unit responded. Selectivity was broadly and roughly evenly distributed, except that many of the 1,389 IC units did not respond to any of the 60 dB stimuli.

**Figure 6.**
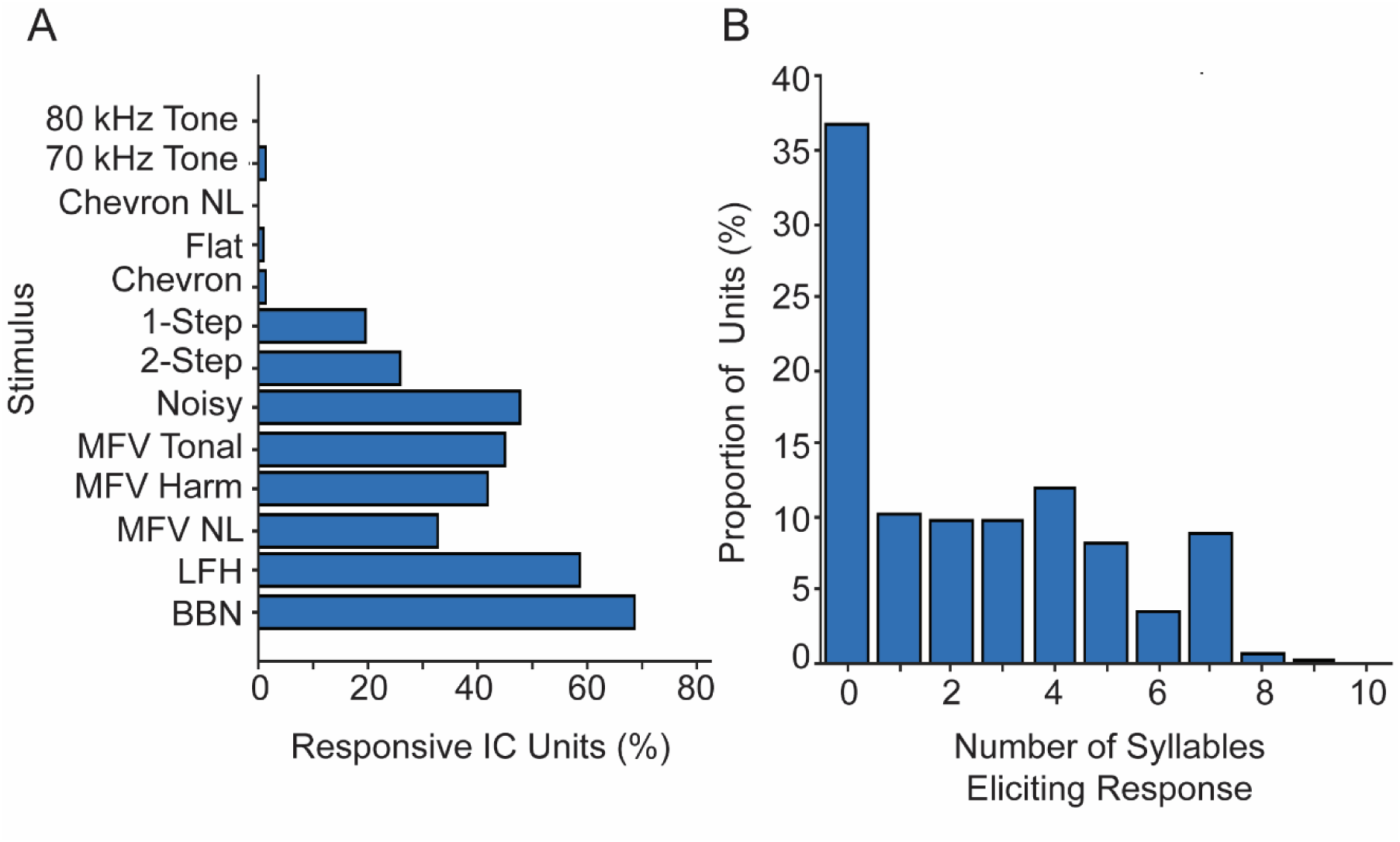
Features of IC population responses to all WAV stimuli. **A.** Comparison of population responses to each WAV stimulus at 60 dB SPL, among IC sound-responsive units (n = 1,212 units). **B.** Selectivity to 10 vocal stimuli among all IC units (n = 1,389 units).

The example units in Figure 4 suggest that features of the FRA may well predict the population response to mouse vocal stimuli. For example, the population-wide responsiveness to tones above 8 kHz (Fig. 3) appears to explain the response of even high-CF units to stimuli with fundamentals below 20 kHz (Figs. 5B-D). How well does the high frequency limit of FRAs/FRCs explain whether lower-CF units can respond to USVs?

To assess this, we quantified the spectral overlap between the FRC of each unit at 60 dB SPL and the bandwidth of each vocal stimulus, as illustrated in Figure 7A. Figures 7B-F then relate the vocal responses of IC units to the extent of FRC—call overlap, with particular attention to calls with higher frequency fundamentals. This analysis shows that nearly all units responding to a vocalization (*blue histograms*) showed overlap between the FRC and the spectrum of that vocal stimulus. Across the noisy call and USV stimuli, a minority of responses occurred in the absence of FRC—call spectrum overlap. Further, for each USV stimulus (Figs. 7D-F), the percentage of responsive units was higher with increasing spectral overlap, and the percentage of unresponsive units *(red histograms)* was much higher when overlap was small or absent. Finally, the percentage of units responding to a call decreased when the frequency content was too high to allow substantial FRC—call overlap. This is clearly the case with the tonal USVs that evoke responses in a very small number of IC units (Figs. 7E,F).

**Figure 7.**
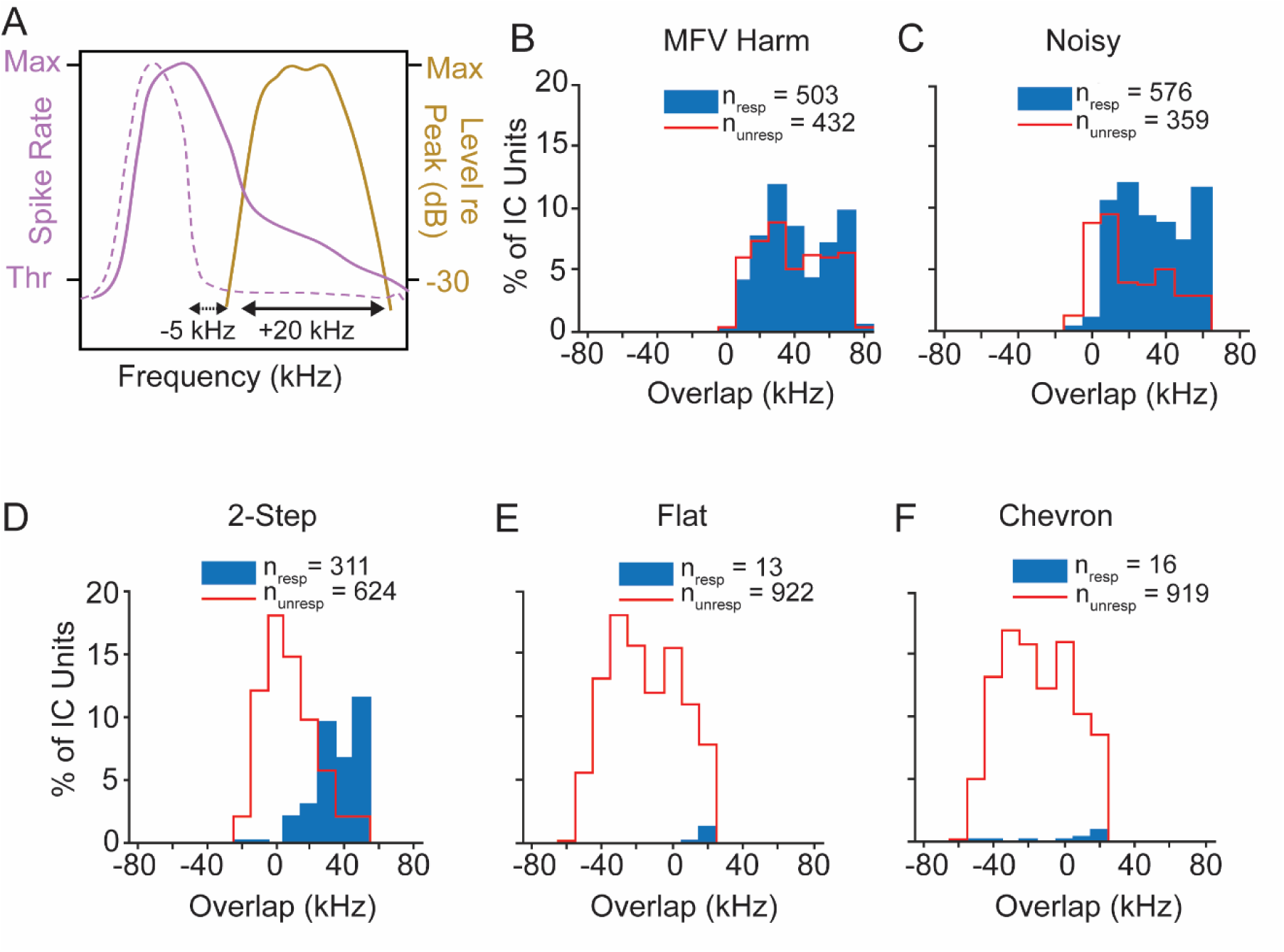
IC units are most likely to respond when call spectrum overlaps with tuning bandwidth. **A.** Schematic showing quantification of overlap between the upper limit of a unit’s frequency response curve at 60 dB SPL (purple lines) and bandwidth of call at -30 dB (gold line). Solid and dashed purple lines show bandwidths of two hypothetical units with (+20 kHz) and without (-5 kHz) overlap between frequency response and call spectrum, respectively. **B-F.** Histograms show extent of overlap among IC units responsive to (filled blue) and non-responsive to (open red) the indicated call at 60 dB SPL peak. Nearly all units that responded to a call showed overlap of frequency tuning with call spectrum. **E-F.** Only a very small percentage of units were responsive to Flat and Chevron calls (< 2%), and most of these showed overlap with the call spectrum. Plots include all units responding to at least one vocal stimulus at 60 dB SPL and having a definable bandwidth of tuning (total n=935 for all stimuli).

We next show that increasing sound level did not alter the close relationship between spectral overlap and responses to USVs. At the peak level of 70 dB SPL, there was little or no change in the number of responsive units to most calls (Figs. 8A,B), but a substantial increase in responsiveness to the chevron call (Figs. 8A-C). Overall, there is little change in the pattern of responsiveness compared to 60 dB SPL stimuli. Further, the increased responses to the higher-level stimuli were mostly among units that displayed substantial FRC-call spectrum overlap (Figs. 8D-F). That is, among units responding to USVs, those without spectral overlap remained a small percentage (<17% for tonal USVs and <6% for stepped USVs).

**Figure 8.**
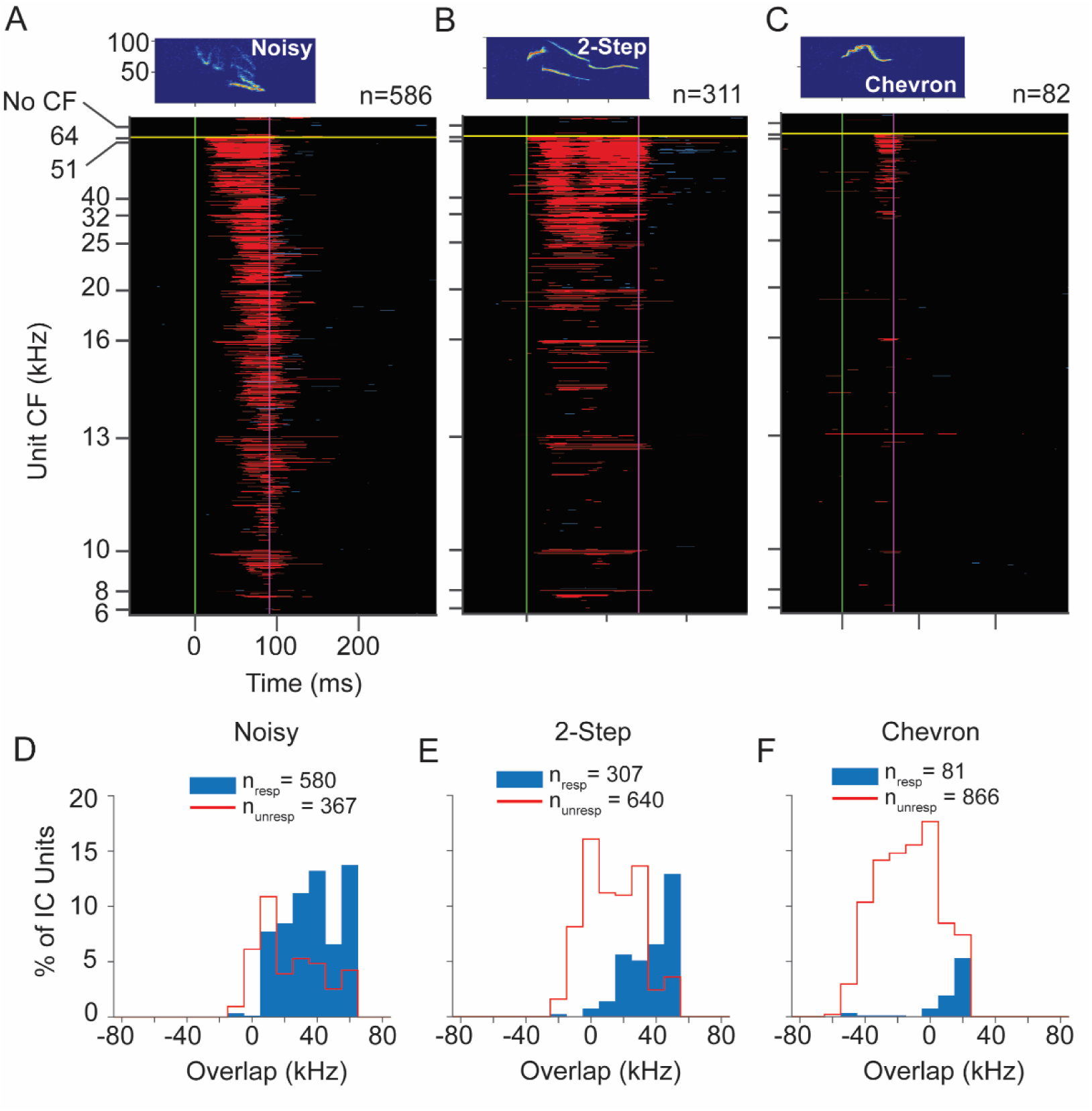
Responses of IC population to WAV stimuli at 70 dB SPL peak. **A-C.** Population responses to the Noisy and 2-Step calls show little or no change in the number of responsive units but there is substantial increase in responsiveness to the chevron call. Overall, there is little change in the pattern of responsiveness compared to 60 dB SPL stimuli. See Figure 5 for protocol. **D-F.** Overlap analysis shows that most units respond to higher frequency vocal signals at 70 dB SPL only when call spectrum overlaps with a unit’s frequency response. See Figure 7 for protocol.

These data indicate that IC unit frequency response properties, as revealed by tonal stimulation, are the primary driver of IC unit responses to mouse vocal stimuli. This analysis also shows that many units did not respond to vocal signals even when spectral overlap occurred. This suggests that additional mechanisms beyond frequency tuning govern selectivity of IC units for social vocalizations.

### Responses to Social Vocalizations: Distributions by Subdivision and Sex

IC subdivisions have different, only partially overlapping projection targets. Our next analysis examined whether these subdivisions represent social vocalizations differently (Fig. 9, Table 9-1). For this analysis, we combined vocalizations into three categories based on spectral content. Category 1 calls have spectral content below 20 kHz (LFH, MFV variants, Noisy). Category 2 includes stepped USVs with energy above 20 kHz and extending above 80 kHz. Category 3 includes USVs with energy exclusively above 60 kHz. Each subdivision displayed the strongest representation of Category 1 vocalizations, moderate representation of Category 2 vocalizations, and minor representation of Category 3 vocalizations. Further, the IC subdivisions similarly represent Category 1 and 3 vocalizations. For the Category 2 calls, the CIC includes a significantly different (greater) representation than in either DCIC or ECIC.

**Figure 9.**
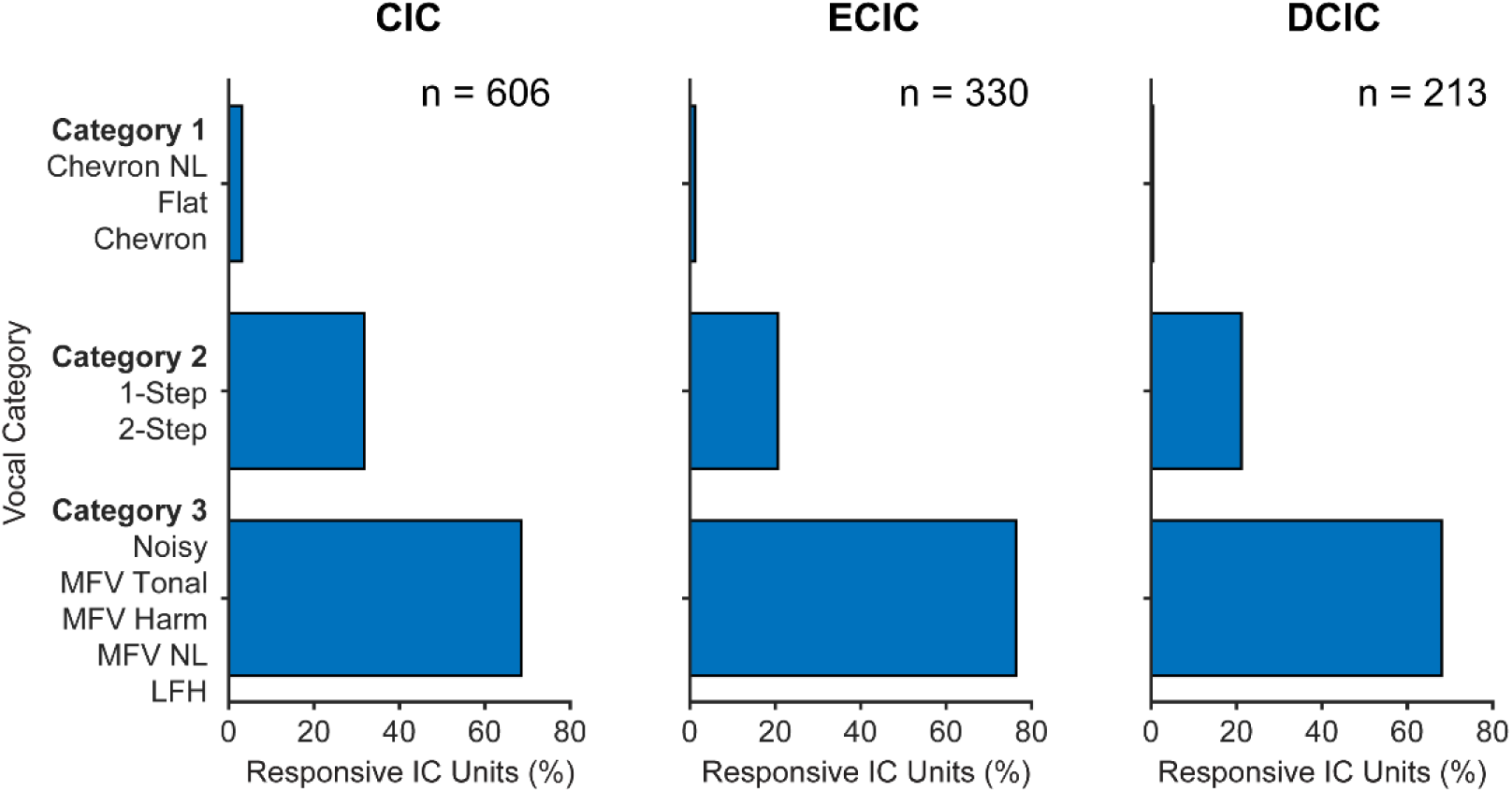
Responsiveness to vocal stimuli as a function of subdivision. **A.** % responsiveness of units to each stimulus is shown by IC subdivision. The broader patterns of responsiveness are similar across subdivisions with some differences in distribution. Overall, category 3 calls evoked the greatest and category 1 the smallest percentage of responses in each subdivision. Category 2 calls evoked a significantly greater percentage of responses within CIC compared to the other subdivisions. Detailed statistics in Table 9-1.

Since we observed sex-based differences in CF distribution and tuning bandwidth in CIC, we examined whether this subdivision displayed sex differences in response to vocal categories. We found no sex differences in the representation of any vocal category (Table 9-1).

### Unanesthetized Recordings

#### Frequency Tuning

In UNANEST mice, 296 IC units had organized FRAs and defined CFs. CF values ranged from 5 to 40 kHz and displayed a broad peak at 10-20 kHz (Fig. 10A). Below 40 kHz, the CF distribution for the UNANEST units closely matched the distribution for ANEST units. There were very few UNANEST units with CFs of 40 kHz and above because these recordings were concentrated in the dorsal IC. For the most sensitive units, there is some evidence that thresholds in UNANEST units were lower than in ANEST units (Fig. 10B). However, our ability to compare thresholds was limited by our minimum sound level for tones (10 dB SPL). It is likely that many units, especially in UNANEST mice, had thresholds below this level (see responses to 70 dB vocalizations).

**Figure 10.**
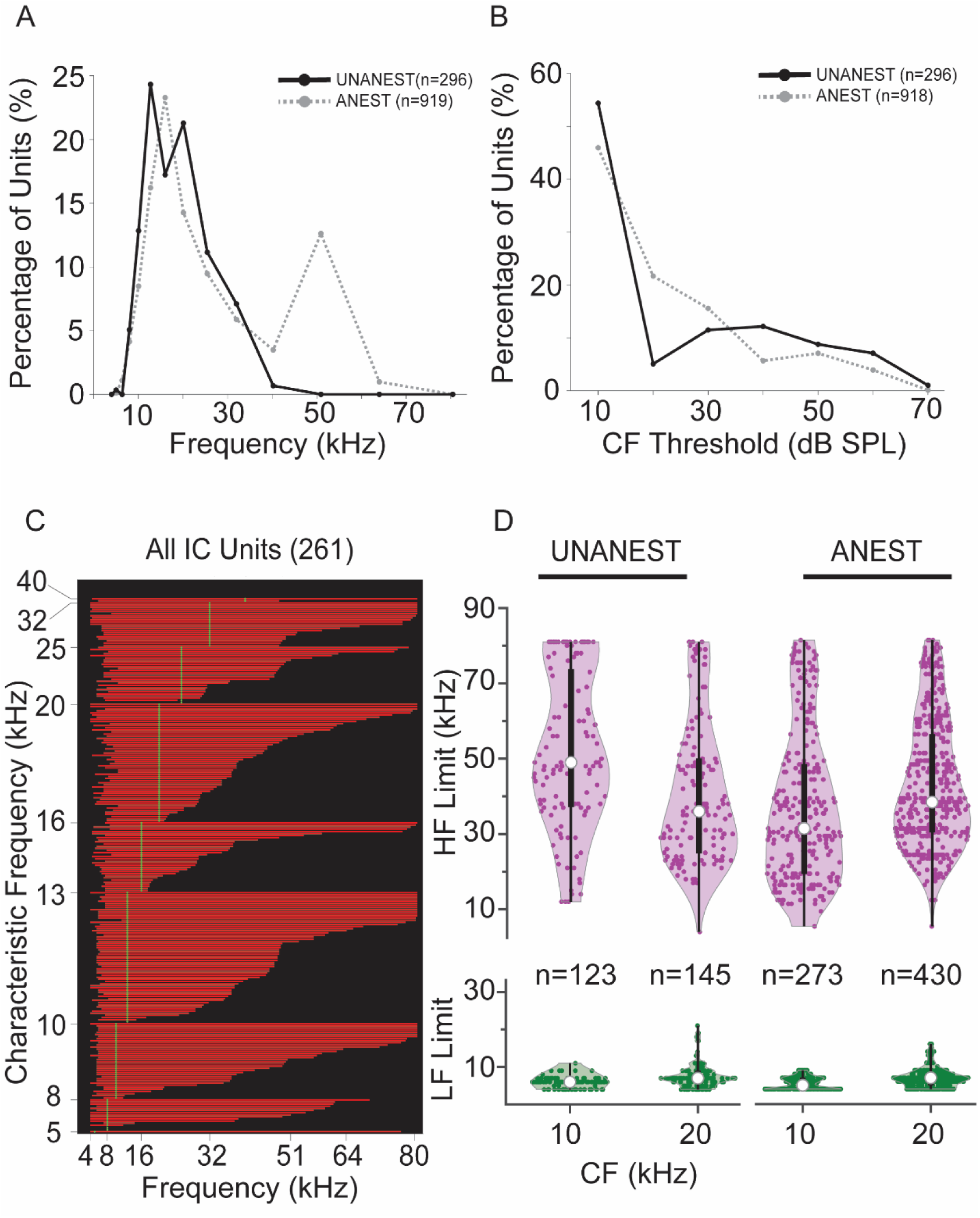
Responses to tonal stimuli in units from UNANEST animals. **A.** Distribution of CFs for UNANEST (black line) and ANEST (dashed gray line) units. The distributions overlap extensively for CFs below 40 kHz, but there were few UNANEST units with CFs above 40 kHz due to the more dorsal recording sites in the UNANEST animals. **B.** Distributions of pure tone thresholds in UNANEST (black line) and ANEST (dashed gray line) units. Since the lowest presented level was 10 dB SPL, many units with 10 dB thresholds may have even lower thresholds. **C.** Frequency response curves displayed as bandwidths (BW) obtained from responses of IC units to tones at 60 dB SPL. Unit BW is displayed as the extent of red line along x-axis (one unit per horizontal line). Units are sorted vertically by CF, then within CF by the upper frequency limit of the BW at 60 dB SPL. BWs varied widely among most CF groups, especially at lower CFs. Compare to ANEST units in Figure 3A. **D.** Comparison of bandwidths for UNANEST (left) and ANEST (right) units. The lower (green) and upper (purple) frequency limits of BWs displayed as violin plots in two octave-wide frequency bands, centered at 10 and 20 kHz. For all violin plots, open circle indicates median, bar indicates interquartile range.

The bandwidths of unit responses to pure tones at 60 dB SPL are shown in Figure 10C. As described for ANEST recordings in Figure 3A, these are vertically sorted by unit CF and then by the upper limits of frequency response curves (FRCs). Similar to the ANEST recordings, UNANEST low-CF units (<40 kHz) displayed comparable lower limits (Fig. 10D, green plots) and large variation in upper limits of bandwidths ranging from 8 to 80 kHz (Fig. 10D, purple plots). The main distinctive feature of bandwidth in the UNANEST units was the distribution of high frequency limits of the FRC for the 10 kHz CF group; the median and interquartile range values of the UNANEST units were substantially higher than those in ANEST units (Fig. 10D).

#### Responses to Social Vocalizations

Because the main focus of UNANEST recordings was to examine the responses of low-CF units to USVs, we present in Figures 11A-C the population responses (n = 261 units) to three USVs (2-Step, Chevron, and Flat calls) at peak sound level of 60 dB SPL. The 2-Step USV evoked responses across the range of CFs; the greatest response was among higher-CF units (>16 kHz) but there were several units with low CFs (Fig. 11A). The population response showed temporal structure that appeared to correspond to the early, middle, and late elements of the syllable. Responses to Chevron and Flat USVs were many fewer (Figs. 11B,C). These primarily occurred among units with the highest upper limits of bandwidths and those with CFs >20 kHz. Like the ANEST units (Fig. 5), the UNANEST population response to these USVs was biased toward units with higher CF or higher upper limits in tuning curves, even though the UANEST group did not include any units with CFs above 40 kHz. Population responses for other vocal stimuli are displayed in Fig. 11-1. These show broad responsiveness across the CF range to the non-USV syllables and many fewer responses to tonal USVs, again similar to results from ANEST animals.

**Figure 11.**
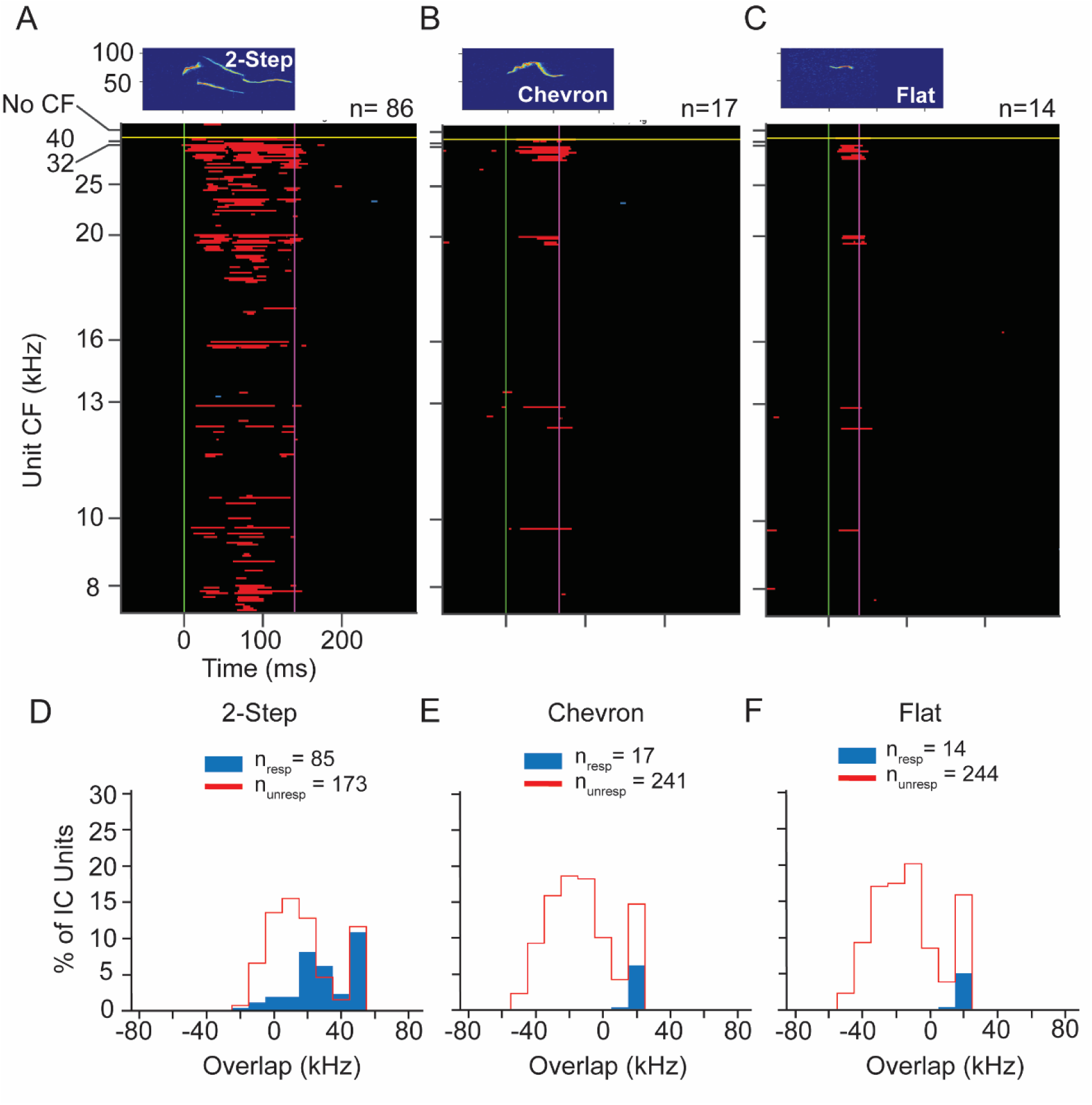
Responses of IC units to WAV stimuli at 60 dB SPL peak in UNANEST mice. **A-C.** Population responses to the 2-Step, Chevron, and Flat calls span a broad range of CFs, but are most common in units with higher CFs. Note that all CFs in UNANEST units were 40 kHz or below, the result of our focus on low frequency units in UNANEST recordings. Each plot shows the same 261 units, the subset of sound-responsive IC units in UNANEST mice that responded to at least one of the 60 dB SPL WAV stimuli. These units are also illustrated in Fig. 10C. See population responses to other 60 dB WAV stimuli in Figure 11-1. **D-F.** Overlap analysis shows that most units respond to higher frequency vocal signals at 60 dB SPL only when call spectrum overlaps with a unit’s frequency response. See Figures 5 and 7 for protocols.

Next, we assessed the degree to which the upper frequency limits of unit FRCs can predict responsiveness to USVs in UNANEST animals. We used the same analysis as described previously for ANEST recordings by quantifying the spectral overlap between the unit FRC and each vocalization. Figures 11D-F shows the results of this analysis for the 2-Step, Chevron, and Flat USVs. For the 2-Step call (Fig. 11D), most responding units showed extensive overlap between the FRC and call spectrum, but a small number of units showed minimal or no overlap between these. Moreover, the percentage of units responding to this syllable was much higher when its FRC overlapped extensively with the syllable bandwidth (e.g., at 50 kHz). For the Chevron and Flat calls, all responding units showed extensive overlap between the FRC and call spectrum (Figs. 11E,F). These results correspond closely to the results for ANEST units.

Analysis of responses to vocalizations presented at 70 dB SPL peak was complicated by the apparently lower thresholds of many UNANEST units (Fig. 10B). At this sound level, we found that many UNANEST units responded to the background noise within the WAV files that preceded the vocal stimulus. Such units (n = 86) were removed from the analysis. As a result, the number of analyzed units (n = 180 units) is much less for 70 dB SPL stimuli, especially at CFs of 32 and 40 kHz. These population response plots are thus difficult to compare with those in Figures 5, 8, and 11. For the 2-Step call, units responded across most of the CF range, emphasizing higher CFs (Fig. 12A). Responses to Chevron and Flat calls were few. Units responding to the Chevron syllable were almost all below 20 kHz (Fig. 12B) and all responses to the Flat syllable were by units with CF of 16 kHz and above (Fig. 12C).

**Figure 12.**
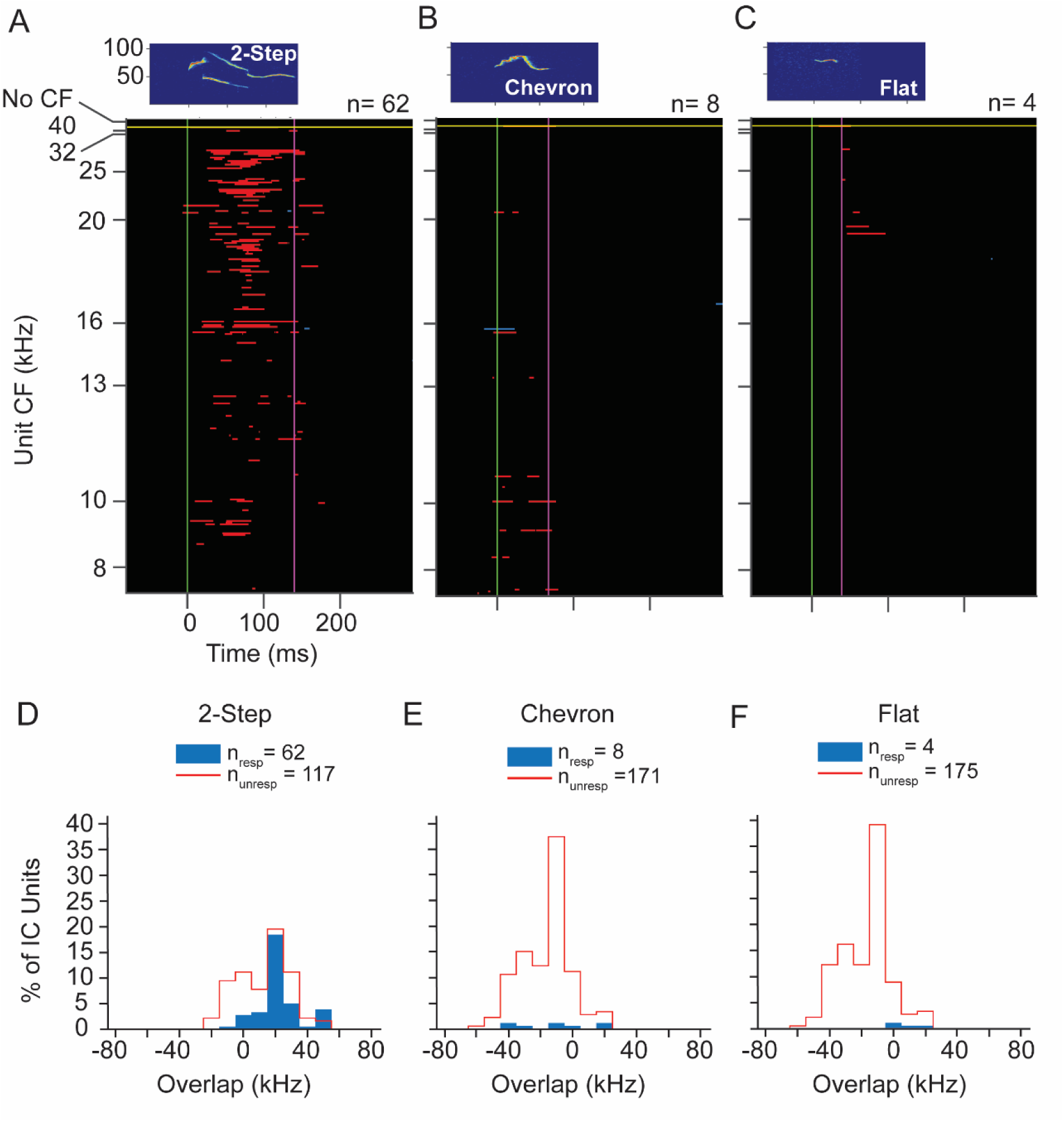
Responses of IC units to WAV stimuli at 70 dB SPL peak in UNANEST mice. **A-C.** Population responses to the 2-Step, Chevron, and Flat calls. As in Figure 11A-C, all CFs in UNANEST units were 40 kHz or below. The 70 dB analysis includes many fewer units than at 60 dB, especially at higher CFs, due to removal of units that responded to the noise background of WAV stimuli. Each plot shows the same 266 units, the subset of sound-responsive IC units in UNANEST mice that responded to at least one of the 70 dB SPL WAV stimuli. Of these, 180 units showed valid responses that were analyzed further, while 86 units were removed from further analyses. **D-F.** Overlap analysis shows that most units respond to 2-Step USV at 70 dB SPL only when call spectrum overlaps with a unit’s frequency response curve. For all three USVs, some units responded even when there was no overlap between call spectrum and their frequency response curves. See Figures 5 and 7 for protocols.

The overlap analysis, limited by lower numbers of higher-CF units, nonetheless shows two noteworthy features (Figs. 12D-F). First, for the 2-Step syllable, units with extensive overlap formed the largest number of responsive units and the highest likelihood of response (Fig. 12D). Second, there were small numbers of units responding to each syllable that displayed no overlap, with the upper limit of their tuning bandwidth as much as 40 dB below the lower frequency limit of the syllable spectrum. This result corresponds closely to 70 dB SPL results in the ANEST animals (Fig. 8).

## DISCUSSION

An overall goal of this study was to place the responses to ultrasonic vocalizations (USVs) in the mouse IC within the broader context of IC frequency tuning and responses to a wider array of social vocalizations. We describe responses of IC neurons to several categories of mouse social vocalizations, relating these responses to frequency tuning, IC subdivisions, and the sex of experimental animals. Overall, we found that responses to lower frequency, mostly broadband social vocalizations were widespread in IC, well represented throughout the tonotopic axis, across subdivisions, and in both females and males. In contrast, responses to USVs were more limited across all subdivisions and both sexes. For most units, responses to vocal signals occurred only when a unit’s frequency response area(s) overlapped with the spectra of the vocal signals. Since tuning to the frequencies contained within the highest frequency USVs is limited in IC, the response to these USVs is correspondingly limited. These results highlight a paradox of USV processing in rodent auditory systems: although USVs are the most abundant social vocalization, their representation and the representation of corresponding frequencies is less than those of the lower frequency social vocalizations.

### Substrates for vocal responses in IC: frequency tuning

Sensory neuroscientific studies generally, and neuroethological studies specifically, have shown that the behavioral significance of sensory signals is often matched by expanded representations of sensory surfaces and central sensory brain structures that analyze these signals. In hearing, this has been well demonstrated by the overrepresentation of biosonar-related frequencies in bat peripheral (Kössl and Vater, 1985; Vater et al., 1985) and central auditory structures (Suga and Jen, 1976; Schuller and Pollak, 1979; Ostwald, 1984; Zook et al., 1985). It is thus perplexing that in mouse and rat auditory systems, representation of frequencies corresponding to ethologically important USVs appears to be so limited (Portfors, 2018). For example, in the IC of mice and rats, none or very few neurons have CFs at the peak frequency of many USVs (>60 kHz in mice; >40 kHz in rats) (Stiebler and Ehret, 1985; Egorova et al., 2001; Portfors and Felix, 2005; Malmierca et al., 2008).

With respect to the mouse IC, two hypotheses have been advanced to explain this apparent mismatch. Portfors and colleagues have identified responses to USVs among low frequency tuned neurons, consistent with the generation of cochlear distortion products by some USVs (Portfors et al., 2009; Portfors and Roberts, 2014; Portfors, 2018). More recently, Garcia-Lazaro and colleagues (2015) have reported that the representation of ultrasonic frequencies in IC is in fact expanded compared to the auditory periphery or lower brainstem, allowing for substantial responsiveness to mouse USVs in the IC. Here, we evaluate the frequency representation in IC and later consider both hypotheses in light of our results on population responses to vocal signals.

Our recordings sampled extensively throughout the IC and documented recording site location relative to an existing atlas (Franklin and Paxinos, 2007). While the distribution of recording sites contained gaps, we sampled most of the rostrocaudal and mediolateral extent of the central nucleus, and all of the dorsoventral extent. There are some differences in coverage of the dorsal and external IC between females and males, but the combined coverage was extensive within the caudal 60% of the IC that was accessible in dorso-ventral penetrations. Within this framework, we show that the distribution of characteristic frequency (CF) generally conformed to previous studies in the mouse IC, with the majority of CFs below 40 kHz (Stiebler and Ehret, 1985; Egorova et al, 2001, Portfors and Felix, 2005). Our data in CBA/CaJ mice show the peak of the CF distribution in the 10-20 kHz range, both for the IC as a whole and for each subdivision. However, we found an additional, substantial peak at ∼50 kHz that was particularly robust in the male CIC. Some previous studies have documented aspects of this higher frequency peak (Portfors and Felix, 2005; Mayko et al., 2012; Garcia-Lazaro et al., 2015).

The study by Garcia-Lazaro and colleagues (2015) reported that the IC distribution of CFs in CBA/CaJ mice (the same strain as we used) heavily emphasizes frequencies above 40 kHz. Further, by comparison to cochlear responses and representation in the cochlear nucleus, they report that the IC frequency representation shows disproportionately large representation of ultrasonic frequencies. Our data support neither of these conclusions. We found that units with CFs of 40 kHz and above constituted no more than 15% of the overall IC and 20% of CIC populations. Further, these distributions did not appear to be substantially different from the CF distribution in auditory nerve fibers (Taberner and Liberman, 2005). Consequently, we believe that the paradox of USV processing in the mouse IC remains: CFs of units are substantially below the frequencies of the most abundant category of social vocalizations in these animals.

In the IC, as elsewhere in the auditory system, the CF of a neuron is a poor predictor of its responses to vocalizations (Leroy & Wenstrup, 2000; Klug et al., 2002; Holmstrom et al., 2007; Portfors et al., 2009; Mayko et al., 2012, Portfors, 2018). As a result, we analyzed the width of frequency tuning functions to understand how neurons might respond to vocalizations with energy restricted to bands away from the CF. There were two noteworthy results. First, at 60 dB SPL, nearly all units responded to tones as low as 8-10 kHz, suggesting that most high-CF neurons would respond to the low frequency, non-USV vocal signals. This was likely due to the low frequency tails of higher CF tuning curves. Second, neurons with low-CFs displayed a broad range of upper limits to their tuning curves, so that even some units with CFs of 8 kHz responded to tones above 64 kHz. This could result either from a single broad frequency tuning curve or from secondary tuning peaks (Egorova et al., 2001; Portfors and Felix, 2005). This suggests mechanisms by which low frequency units respond to high frequency USVs, and at least a partial solution to the paradox of USV processing in the mouse IC. Across subdivisions, the high frequency limits of IC neurons could allow processing and responses to many types of USVs, especially at higher sound levels, even if these subdivisions have few high-CF units.

### Responses of IC neurons to social vocalizations

Adult mice emit several categories of social vocalizations within a range of behavioral and social contexts. USVs are the most common category, emitted by males and sometimes females during mating and by both males and females in other social contexts (Maggio et al., 1983; Maggio and Whitney, 1985; Scattoni et al., 2009; Neunuebel et al., 2015; Grimsley et al., 2016). USVs may be broadly categorized as tonal, with continuous frequency-time functions, or stepped, with abrupt frequency transitions. They may also include harmonics, either a higher harmonic with energy above 80-100 kHz, or a fundamental, in the 30-40 kHz range, to the dominant second harmonic. USVs in courtship and mating show more steps, more complex types, more harmonics, and an overall shift to lower minimum frequencies as the interaction progresses or its intensity increases (Gaub et al., 2016; Keesom and Hurley, 2016; Matsumoto and Okanoya, 2016; ; Ghasemahmad et al., 2022). Further, the distribution of USV peak frequencies shifts to lower frequencies in several contexts: by males in the presence of females (Hanson and Hurley, 2012), in aversive contexts such as restraint or isolation (Grimsley et al., 2016). These considerations suggest that, for adult USVs, lower frequency content may signal increasing emotional intensity of the sender.

Our data show that many more IC units responded to USVs with lower frequency content. At a peak sound level of 60 dB SPL, about one-quarter of IC units responded to the stepped calls, less than 2% responded to flat and chevron USVs with energy only above 60 kHz, and two units responded to the non-linear chevron syllable with energy limited to 80 kHz and above. Playback of these syllables at a higher sound level (70 dB SPL peak) results in somewhat larger numbers of responses, but no change to these fundamental patterns of population response. Thus, behaviors that elicit more USV syllables with fundamentals in the 30-40 kHz range (as occurs during mating) and other USVs with lower minimum frequencies will excite many more IC neurons. In our study, IC auditory responses are better matched to the lower frequency components of USVs that signal heightened emotional intensity.

At least three other categories of mouse vocalizations have energy below 20 kHz. The Low Frequency Harmonic call (Grimsley et al., 2011), i.e., the mouse squeak, has its fundamental near 5 kHz and multiple harmonics extending well into the ultrasonic range. It is emitted by females during mating (Sewell, 1972; Grimsley et al., 2013, Keesom and Hurley, 2016) and by both sexes during behavioral contexts associated with threat or pain (Gourbal et al., 2004; Williams et al., 2008; Grimsley et al., 2016). Several studies have established the salience of this call (Grimsley et al., 2013; Keesom and Hurley, 2016; Niemczura et al., 2020). Sixty percent of IC units responded to our exemplar. While the duration and energy within the ultrasonic range can vary substantially in some contexts (Grimsley et al., 2016; Finton et al., 2017), these variations may be encoded by the timing of responses, which seem to vary with the duration of the spectral components in the calls. Recent work shows both temporal patterning to LFH calls and selectivity to different variants (Polese et al., 2021). Strong responsiveness to LFH calls is maintained in at least one indirect target of the IC, the basolateral amygdala (Grimsley et al., 2013).

Mid-frequency vocalizations (MFVs), more recently characterized (Grimsley et al., 2016), occur commonly when mice are restrained. They are increasingly emitted with activation of a noradrenergic pathway to the rostromedial tegmental nucleus associated with aversive behavior (Dornellas et al., 2021). In listening mice, the calls are stressful (Niemczura et al., 2020) and evoke defensive responses and altered neuromodulator release into the amygdala that suggest increased vigilance and reduced reward (Ghasemahmad et al., 2022). MFVs vary in duration, harmonic content, and non-linear features (Grimsley et al., 2016), but most have fundamentals in the 10-18 kHz range. Depending on MFV type, this call category evokes responses in 33-42% of IC units, throughout the tonotopic range.

Noisy calls, with both frequency modulations and chaotic elements, often have energy below 20 kHz but extend well into the ultrasound range. They have been most associated with isolation (Grimsley et al, 2016) and their playback results in increased plasma corticosterone (Niemczura et al., 2020). However, the function of this call type remains poorly understood. Nonetheless, with broad spectral content, the Noisy call excites nearly half of IC neurons (48%) throughout the tonotopic range with good preservation of spectrotemporal information within the call.

While our use of multi-unit responses has the potential to overestimate the responsiveness of individual neurons and to underestimate both potential response selectivity and inter-neuron variability, our approach permits a few conclusions about responsiveness and selectivity. First, high responsiveness (and low selectivity) is primarily to the lower frequency and broadband stimuli. Across IC, responsiveness to LFH, MFV, and Noisy syllables averaged 45% of the IC sample. More specifically, when there was overlap between the spectra of these syllables and tuning width of units, over half of units responded (see Fig. 7B,C). In contrast, responsiveness was much lower and selectivity much higher in neuronal responses to USVs, even when there was overlap between syllable spectra and tuning bandwidth of units (Fig. 7D-F). The mechanisms of selectivity for USVs appear to depend on multi-peaked tuning, combination sensitivity, cochlear non-linearities, and inhibition (Portfors et al., 2009; Mayko et al., 2012; Portfors, 2018). Mechanisms of selectivity may further depend on context-dependent modulation of IC responses, e.g., by serotonin (Hurley & Pollak, 2005; Petersen & Hurley, 2017; Polese et al., 2021).

Few studies across mammals and other vertebrates have examined how vocal responses are distributed across IC subdivisions (Aitkin et al., 1994; Šuta et al., 2003). These studies in cat and guinea pig show relatively similar high levels of responsiveness to vocalizations across subdivisions. Based on our comparison of frequency representation, tuning properties, and vocal responses across IC subdivisions, we conclude that each subdivision and its projection target(s) in the auditory thalamus will respond in similar proportions to the mouse vocal repertoire. Thus, all subdivisions will convey information about the three categories of broadband and lower frequency vocalizations. The main difference among subdivisions concerns their population responses to USVs. The CIC population, and by inference its targets in the ventral MG, shows greater responsiveness to all USVs, and is likely to respond better to the higher frequency USVs due to the enhanced numbers of neurons with both higher CFs and higher upper frequency limit. Overall, information about USVs that ascends to the auditory thalamus appears to be biased to the lower frequencies within these calls associated with increased emotional intensity.

### Mechanisms of IC responses to USVs

Here we use our data to evaluate previous proposals to explain responses to USVs in the mouse auditory system.

Portfors and colleagues report extensive responsiveness to USVs among low-CF units in the mouse IC. They propose that such responses depend on low frequency cochlear distortion products created by combinations of very high frequency USV components or by the frequency modulations in USVs (Portfors et al., 2009: Portfors and Roberts, 2014; Portfors, 2018). Our data do not well support cochlear distortion products as the major explanation of USV responsiveness, at least for stimuli presented at 60 or 70 dB SPL. By comparing USV responses with each unit’s tuning width, we found that almost all USV-responsive neurons displayed spectral overlap. We recognized that our overlap analysis has some limitations: for example, many spectral components of the vocalizations occur at levels below the peak playback level (60 or 70 dB), whereas the frequency response curve is obtained at precisely 60 or 70 dB SPL. This would likely result is some over-estimate of overlap. A more precise method would incorporate more information about the amplitude spectrum of calls and unit FRAs. This may increase the proportion of units that are responsive but without spectral overlap, but it is unlikely to change the major conclusion. That is because most units responding to USVs (except in UNANEST 70 dB data) showed bandwidths that overlapped with call spectrum by 20 kHz or more.

For most IC neurons, therefore, the features of the pure tone FRA, especially its bandwidth, are sufficient to explain responsiveness to USVs. Specifically, less than 20% of USV responses occur without spectral overlap. These require other explanations, such as the cochlear distortion product hypothesis of Portfors and colleagues. These considerations lead us to conclude that cochlear distortions are a relatively minor contribution to IC vocal responses. A caveat to this conclusion is that distortion products may lead to IC excitatory responses mainly at sound levels higher than 70 dB. Indeed, the stimuli used in studies by Portfors and colleagues were often at higher sound levels. However, these higher-level stimuli would also interact with neurons’ broader tuning bandwidth at the higher level.

We were concerned that our use of urethane anesthesia altered cochlear sensitivity, tuning, or non-linearities in a way that could affect IC responsiveness to vocalizations. Previous work suggests that urethane anesthesia may not affect the amplitude of distortion product otoacoustic emissions (DPOAEs), but clearly reduces the contralaterally evoked suppression of the DPOAE (Chambers et al., 2012). In our study, we saw no evidence that anesthetic condition affects our conclusion that cochlear distortions form a relatively minor contribution to IC vocal responses.

The alternative explanation of USV responses was proposed by Garcia-Lazaro and colleagues (2015): that there are many USV responses in the IC and that these emanate from high-CF neurons that other studies missed. We disagree with these conclusions for two reasons. First, as noted above, we find no evidence that the IC contains larger numbers of high-CF units than low-CF units, despite our attempts to sample extensively in the rostro-caudal, medio-lateral, and dorso-ventral dimensions of the IC. For example, units tuned ≥ 60 kHz were encountered within a single penetration of the 4-shank electrode, out of 23 animals. These units occur but are likely rare. Second, although the IC contains a significant population of units tuned near 50 kHz, many of these units are not well responsive to the tonal USVs that we tested with energy only above 64 kHz. If the explanation for USV responses is that there are large numbers of high-CF units, we see neither the large numbers of high-CF units nor the responsiveness to tonal USVs expected of such units.

For these reasons, we suggest that the dominant form of responsiveness to USVs depends on basic features of unit frequency tuning, and that these features emphasize responses to the lower frequencies (30-60 kHz) within USVs. Further, we propose that these are frequencies that signal intense emotions in both mating and other behavioral contexts. Of course, mice do respond to higher frequency USVs, but our data suggest that such responses depend on high sound levels, either to engage the broader FRA bandwidth at high sound levels or to generate cochlear distortion products.

### Vocal responses in auditory cortex

How does this representation of the mouse vocal repertoire, especially USVs, compare to auditory cortex? The major substrate of USV responses—cortical neurons tuned above 45 kHz—was originally thought to be located within an ultrasonic field (UF) outside of the tonotopically organized primary (AI) or anterior (AAF) auditory fields (Stiebler et al., 1997). More recent studies have revised this to consider that neurons with ultrasonic CFs >45 kHz are part of several auditory fields, including AI, AAF, and a dorsomedial field (DM) (Guo et al., 2012; Issa et al., 2014; Tsukano et al., 2015). DM, a projection target of the ventral division of the medial geniculate body, is particularly responsive to ultrasonic frequencies (Tsukano et al., 2015). Overall, however, it remains unclear whether individual auditory cortical fields or auditory cortex overall reflect the distribution of ultrasonic frequencies observed in the IC or its subdivisions, or whether there is an expansion of these frequencies in auditory cortex as has been described in the rat (Kim & Bao, 2013).

Studies of vocal responses in mouse auditory cortex have mostly focused on pup vocalizations within auditory cortical regions containing high-CF neurons (Liu & Schreiner, 2007; Galindo-Leon et al., 2009; Shepard et al., 2015). These studies report many responses to vocalizations with spectra above 50 kHz, but it is unclear how common are these responses in AC generally or among high-CF neurons. Responses to adult USVs have been rarely explored in auditory cortex (Shepard et al., 2015; Tsukano et al., 2015). Tsukano and colleagues report broad excitation in DM and AI resulting from male USVs with peak energy above 60kHz, but inspection of their stimuli show several USVs with fundamentals in the 35-40 kHz range that may have contributed to responses (Fig. 14; (Tsukano et al., 2015). We are not aware of neurophysiological studies that have examined responses to the non-USV syllables tested here. As a result, the overall AC representation of social vocalizations is not understood. We propose that it follows the general principles observed in the IC: strong representation of non-USV calls, significant representation of USVs with lower ultrasonic frequency content, and restricted representation of USVs with spectral content only above 60 kHz that is mostly limited to higher sound levels.

### Sex Differences in the Mouse IC

This study utilized comparable numbers of female and male subjects to test whether auditory midbrain responses to simple acoustic stimuli and vocalizations display sex or estrous differences. Our motivations were both general (sex differences in vocal and reproductive behaviors) and specific (sex and estrous differences in behavioral, hormonal, and neurochemical responses to playback of USVs or vocal sequences in mating that combined USVs and LFH calls (Niemczura et al., 2020; Ghasemahmad et al., 2022). We sought to test whether these different responses to vocal playback might have a basis in auditory sensitivity within the ascending auditory pathway.

Our analysis revealed statistically significant differences in the distribution of CFs between females and males, with males showing a larger proportion of units with CFs near 50 kHz. This result is particularly strong in the CIC. Also in CIC, units with CFs in an octave band centered at 20 kHz had higher-frequency cutoffs in females than in males. These female/male differences may be offsetting: females had fewer of the highest CFs but female units with lower CFs (20 kHz band) had higher frequency cutoffs. These offsetting differences may explain why we observed no sex differences in the representation of any category of social vocalization.

Our study also reveals the challenges of analysis of sex-based differences in auditory responses. Due to the heterogeneity of auditory response properties and their distribution throughout IC, a key consideration is whether there is comparable sampling across the IC for females and males, and for females in different estrous stages. As we have shown (Fig. 1), sampling of units in female and male subjects overlapped extensively within CIC, supporting our in-depth analysis of properties within this subdivision. We are less confident of sex- or estrous-based comparisons outside of CIC because uneven sampling throughout ECIC and DCIC could bias these results even more.

Overall, we believe these results are intriguing but preliminary. For estrous state analyses, chronic recordings will be powerful tools to assess such differences. Understanding sex differences will be a greater challenge, since studies must ensure comparable sampling within structures of the auditory system. The IC, with its complex distribution of response properties, poses particular challenges.

## ACKNOWLEDGMENTS

This work was supported by research grant R01DC00937 from the National Institute on Deafness and Other Communication Disorders to J. Wenstrup and Alexander Galazyuk. R. Patel and D. Kazimiersky were supported by Summer Research Fellow Program awards from the NEOMED Office of Research to S. Shanbhag. We thank Debin Lei and Jacob Rose for technical assistance and Zahra Ghasemahmad for providing vocal stimuli.

## EXTENDED FIGURES AND TABLES

**Table 3-1.**
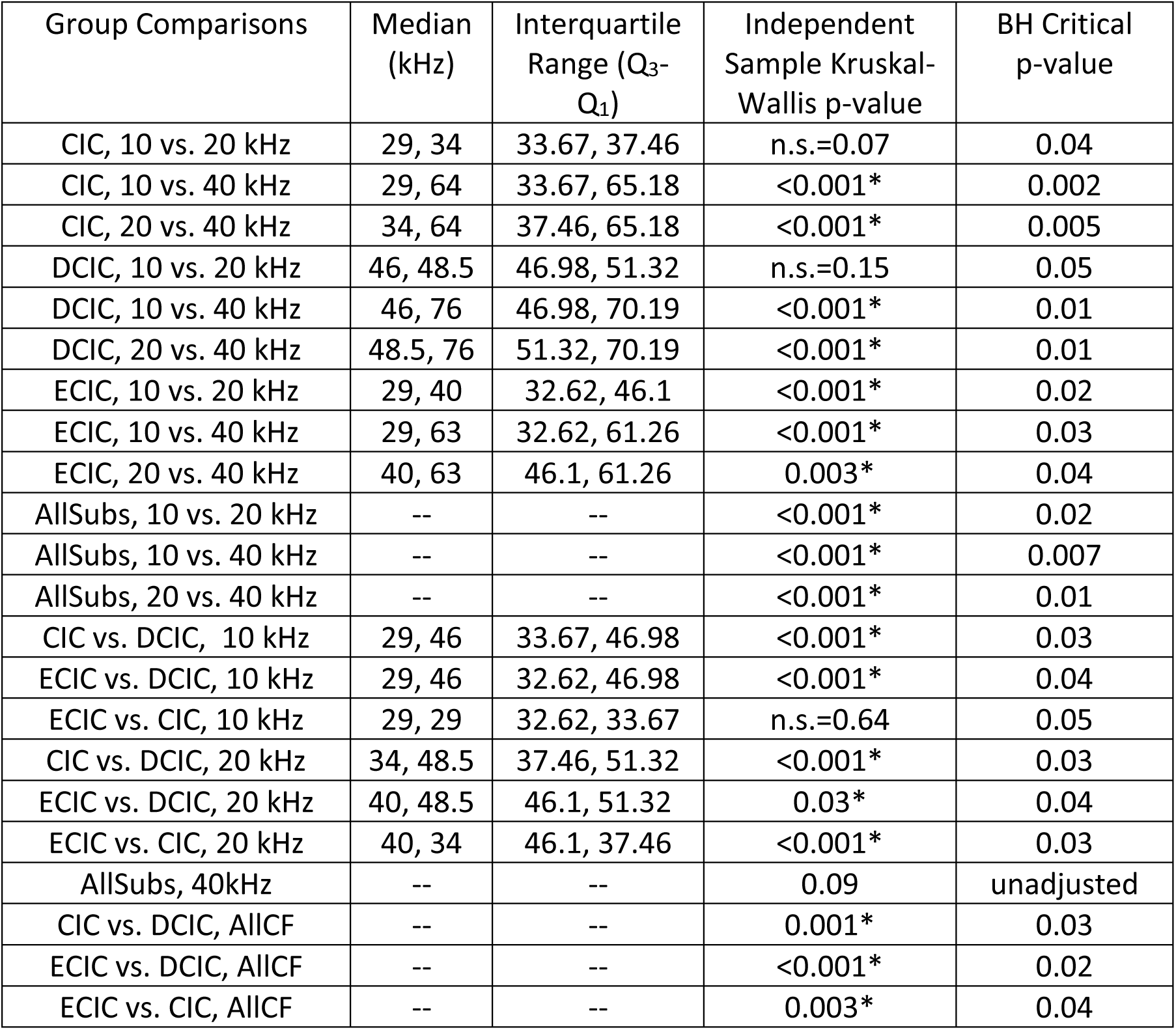
Statistical comparisons of the distributions of upper frequency limits among IC units by CF group and subdivision. All p-values were derived using independent-samples Kruskal-Wallis tests and adjusted using Benjamini-Hochberg corrections. Significance level = 0.05; False Discovery Rate= 0.05. Statistically significant values are shown with an asterisk. Raw p-values were not adjusted if the null hypothesis (no difference between medians) was not rejected. For all analyses with adjusted p values, degrees of freedom = 2. AllCF = all CF groups combined; AllSubs = all subdivisions combined; n.s.=non-significant.

**Table 3-2.** Statistical comparisons of the distributions of upper frequency limits among CIC units by CF, sex and estrous stage. All p-values were derived using independent-samples Kruskal-Wallis tests and adjusted using Benjamini-Hochberg corrections. Significance level = 0.05; False Discovery Rate= 0.05. Statistically significant values are shown with an asterisk. Raw p-values were not adjusted if the null hypothesis (no difference between medians) was not rejected. For all analyses with adjusted p values, degrees of freedom = 2. F_nonest._ = non-estrus female; F_est._ = estrus female; M = Male; n.s.=non-significant.

| Group Comparisons | Median (kHz) | Interquartile Range (Q<sub>3</sub>-Q<sub>1</sub>) | Independent Sample Kruskal-Wallis p-value | BH Critical p-value |
| --- | --- | --- | --- | --- |
| F<sub>nonest.</sub>, 10 vs. 20 kHz | 19, 37 | 31.26, 38.54 | n.s.=0.08 | 0.05 |
| F<sub>nonest.</sub>, 10 vs. 40 kHz | 19, 64 | 31.26, 64.1 | <0.001* | 0.02 |
| F<sub>nonest.</sub>, 20 vs. 40 kHz | 37, 64 | 38.54, 64.1 | <0.001* | 0.02 |
| F<sub>est.</sub>, 10 vs. 20 kHz | 36, 56 | 36.62, 53.2 | 0.002* | 0.03 |
| F<sub>est.</sub>, 10 vs. 40 kHz | 36, 61 | 36.62, 60 | 0.004* | 0.03 |
| F<sub>est.</sub>, 20 vs. 40 kHz | 56, 61 | 53.2, 60 | n.s.=0.35 | 0.04 |
| M, 10 vs. 20 kHz | 29, 31 | 33.76, 34.55 | n.s.=0.61 | 0.05 |
| M, 10 vs. 40 kHz | 29, 65 | 33.76, 65.95 | <0.001* | <0.001 |
| M, 20 vs. 40 kHz | 31, 65 | 34.55, 65.95 | <0.001* | 0.01 |
| AllSex, 10 vs. 20 kHz | -- | -- | n.s.=0.07 | 0.04 |
| AllSex, 10 vs. 40 kHz | -- | -- | <0.001* | 0.01 |
| AllSex, 20 vs. 40 kHz | -- | -- | <0.001* | 0.01 |
| M vs. F<sub>nonest.</sub>, 20 kHz | 31, 37 | 34.55, 38.54 | 0.03* | 0.04 |
| M vs. F<sub>est.</sub>, 20 kHz | 31, 56 | 34.55, 53.2 | <0.001* | 0.02 |
| F<sub>nonest.</sub> vs. F<sub>est.</sub>, 20kHz | 37, 56 | 38.54, 53.2 | <0.001* | 0.03 |
| AllSex, 10kHz | -- | -- | 0.33 | unadjusted |
| AllSex, 40kHz | -- | -- | 0.23 | unadjusted |

**Figure 5-1.**
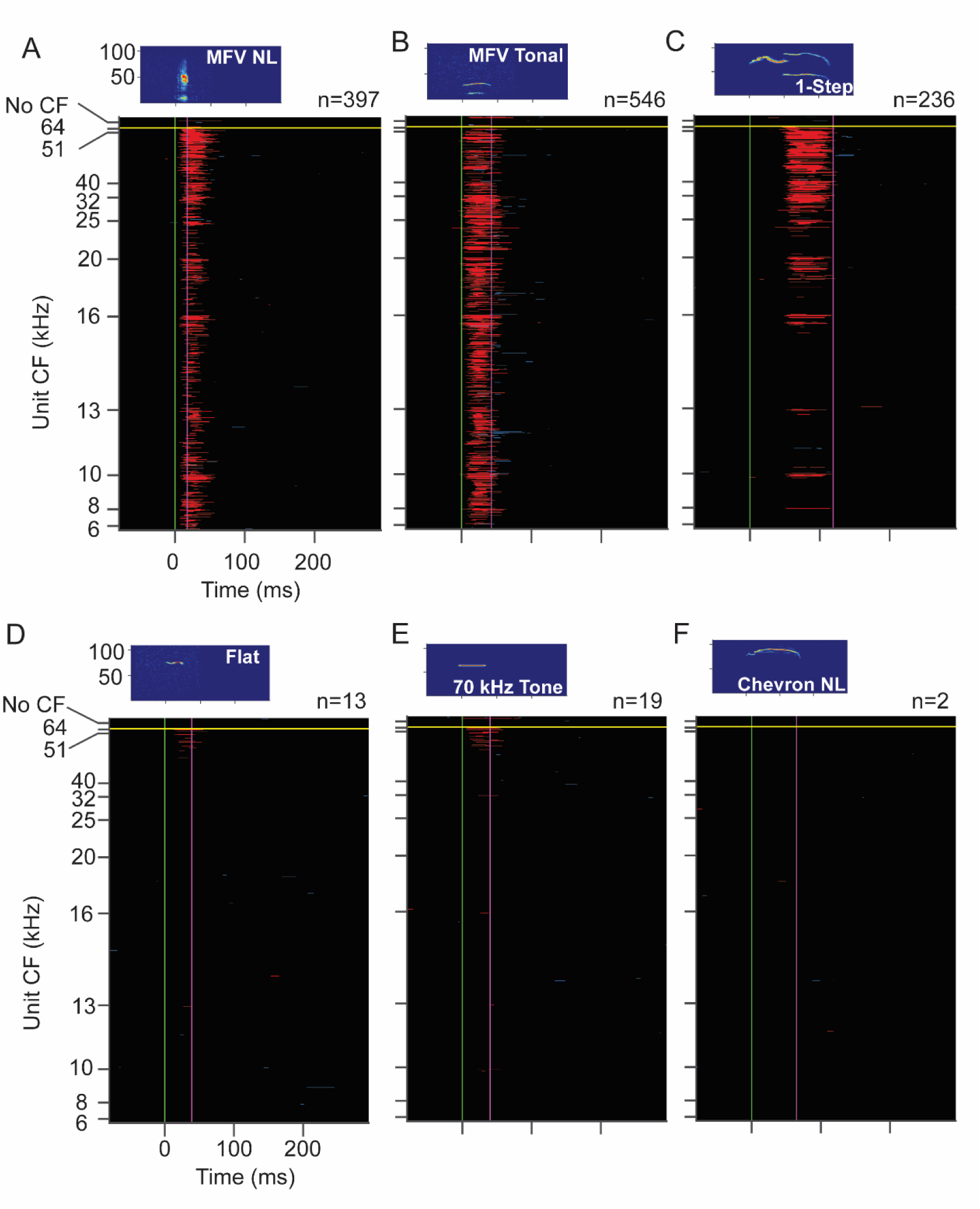
Responses of IC population to WAV stimuli as function of CF. **A-F.** Population response to the WAV stimulus indicated in the sonogram. Each plot shows excitatory (red) and inhibitory (blue) SDF responses to stimuli at 60 dB SPL peak. See Figure 5 for protocol. Each plot shows the same 944 units as in Figures 3A and 5, the subset of 1212 sound-responsive IC units that responded to at least one of the 60 dB SPL WAV stimuli. Stimulus spectrograms and number of responding units shown above main plot.

**Table 9-1.**
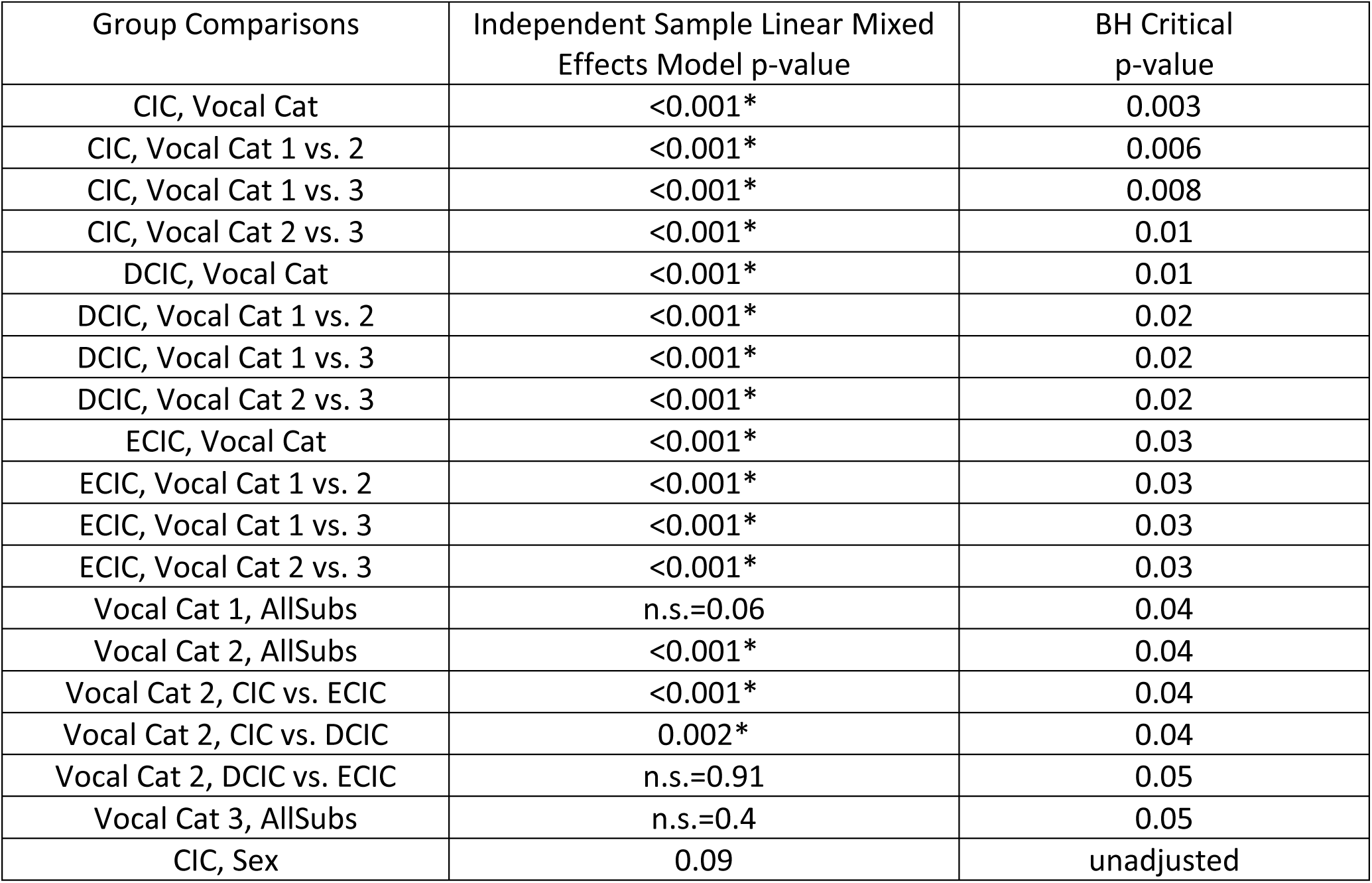
Statistical analyses of IC subdivision responsiveness to categories of social vocalizations. All p-values were derived using a Linear Mixed Effects Model and adjusted using Benjamini-Hochberg corrections. Significance level = 0.05; False Discovery Rate= 0.05. Statistically significant values are shown with an asterisk. There were no sex differences among CIC units in their representation of vocal categories, thus the p-value (0.09) was not adjusted. Vocal Cat = vocal category; AllSubs = all subdivisions combined; n.s.=non-significant.

| Group Comparisons | Independent Sample Linear Mixed Effects Model p-value | BH Critical p-value |
| --- | --- | --- |
| CIC, Vocal Cat | <0.001* | 0.003 |
| CIC, Vocal Cat 1 vs. 2 | <0.001* | 0.006 |
| CIC, Vocal Cat 1 vs. 3 | <0.001* | 0.008 |
| CIC, Vocal Cat 2 vs. 3 | <0.001* | 0.01 |
| DCIC, Vocal Cat | <0.001* | 0.01 |
| DCIC, Vocal Cat 1 vs. 2 | <0.001* | 0.02 |
| DCIC, Vocal Cat 1 vs. 3 | <0.001* | 0.02 |
| DCIC, Vocal Cat 2 vs. 3 | <0.001* | 0.02 |
| ECIC, Vocal Cat | <0.001* | 0.03 |
| ECIC, Vocal Cat 1 vs. 2 | <0.001* | 0.03 |
| ECIC, Vocal Cat 1 vs. 3 | <0.001* | 0.03 |
| ECIC, Vocal Cat 2 vs. 3 | <0.001* | 0.03 |
| Vocal Cat 1, AllSubs | n.s.=0.06 | 0.04 |
| Vocal Cat 2, AllSubs | <0.001* | 0.04 |
| Vocal Cat 2, CIC vs. ECIC | <0.001* | 0.04 |
| Vocal Cat 2, CIC vs. DCIC | 0.002* | 0.04 |
| Vocal Cat 2, DCIC vs. ECIC | n.s.=0.91 | 0.05 |
| Vocal Cat 3, AllSubs | n.s.=0.4 | 0.05 |
| CIC, Sex | 0.09 | unadjusted |

**Figure 11-1.**
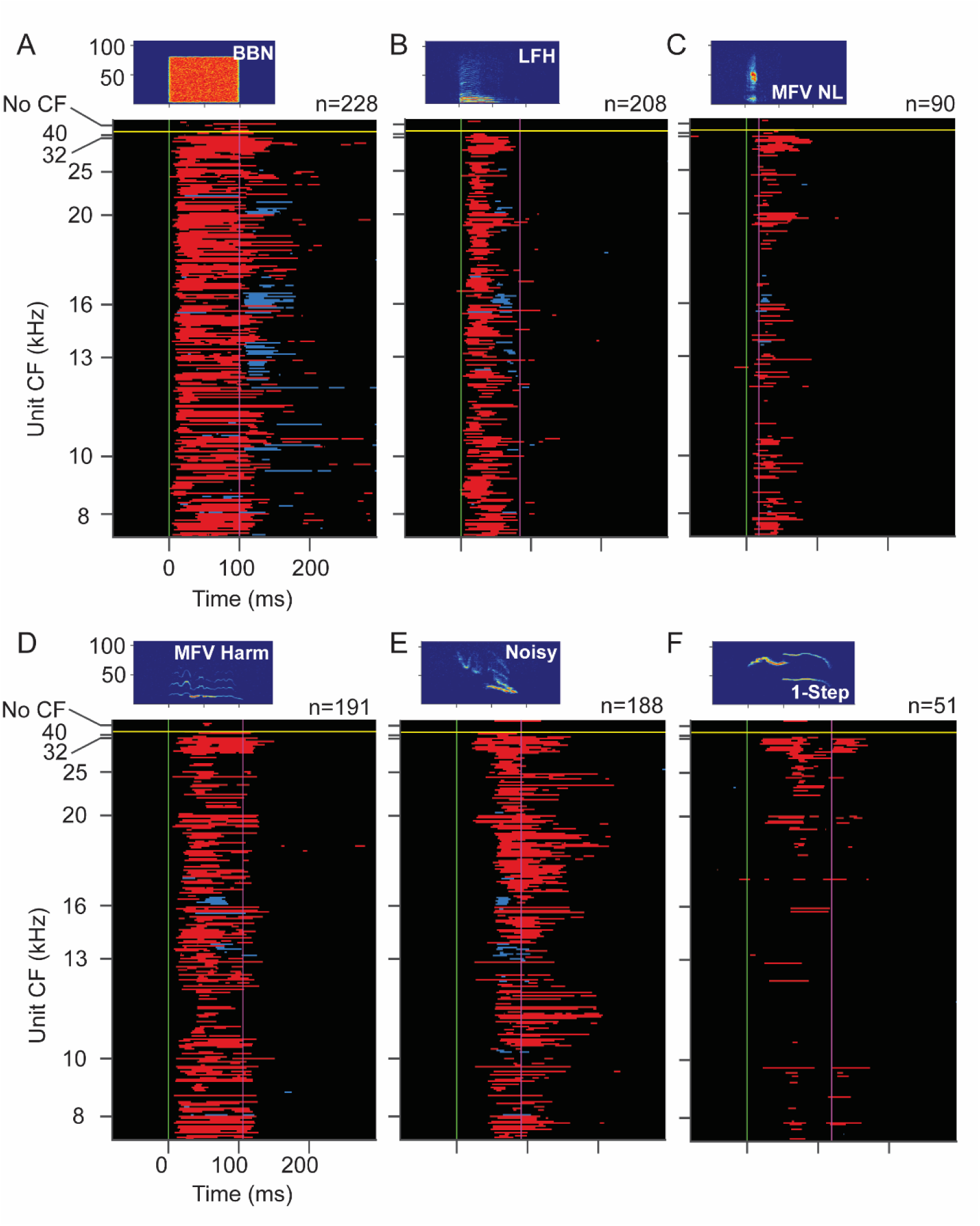
Responses of IC units from UNANEST mice to WAV stimuli as function of CF. **A-F.** Population response to the WAV stimulus indicated in the sonogram. Each plot shows excitatory (red) and inhibitory (blue) SDF responses to stimuli at 60 dB SPL peak. Each horizontal line represents response of one unit. See Figure 5 for protocol. Each plot shows the same 261 units as in Figures 10C and 11, the subset of sound-responsive IC units from UNANEST animals that responded to at least one of the 60 dB SPL WAV stimuli.

## LITERATURE CITED

Aitkin, L., Tran, L., & Syka, J. (1994). The responses of neurons in subdivisions of the inferior colliculus of cats to tonal, noise and vocal stimuli. Experimental Brain Research, 98(1). 10.1007/BF00229109

Bates, D., Mächler, M., Bolker, B. M., & Walker, S. C. (2015). Fitting linear mixed-effects models using lme4. Journal of Statistical Software, 67(1). 10.18637/jss.v067.i01

Brudzynski, S. M. [Ed]. (2018). Handbook of ultrasonic vocalization: A window into the emotional brain. In Handbook of ultrasonic vocalization: A window into the emotional brain.

Chambers, A. R., Hancock, K. E., Maison, S. F., Liberman, M. C., & Polley, D. B. (2012). Sound-evoked olivocochlear activation in unanesthetized mice. JARO - Journal of the Association for Research in Otolaryngology, 13(2). 10.1007/s10162-011-0306-z

Dornellas, A. P. S., Burnham, N. W., Luhn, K. L., Petruzzi, M. V., Thiele, T. E., & Navarro, M. (2021). Activation of locus coeruleus to rostromedial tegmental nucleus (RMTg) noradrenergic pathway blunts binge-like ethanol drinking and induces aversive responses in mice. Neuropharmacology, 199. 10.1016/j.neuropharm.2021.108797

Egorova, M., Ehret, G., Vartanian, I., & Esser, K. H. (2001). Frequency response areas of neurons in the mouse inferior colliculus. I. Threshold and tuning characteristics. Experimental Brain Research, 140(2). 10.1007/s002210100786

Ehret, G. (2005). Infant rodent ultrasounds - A gate to the understanding of sound communication. Behavior Genetics, 35(1). 10.1007/s10519-004-0853-8

Finton, C. J., Keesom, S. M., Hood, K. E., & Hurley, L. M. (2017). What’s in a squeak? Female vocal signals predict the sexual behaviour of male house mice during courtship. Animal Behaviour, 126. 10.1016/j.anbehav.2017.01.021

Franklin, K. B. J., & Paxinos, G. (2007). The Mouse Brain in Stereotaxic Coordinates (map). In Academic Press.

Galindo-Leon, E. E., Lin, F. G., & Liu, R. C. (2009). Inhibitory Plasticity in a Lateral Band Improves Cortical Detection of Natural Vocalizations. Neuron, 62(5). 10.1016/j.neuron.2009.05.001

Garcia-Lazaro, J. A., Shepard, K. N., Miranda, J. A., Liu, R. C., & Lesica, N. A. (2015). An overrepresentation of high frequencies in the mouse inferior colliculus supports the processing of ultrasonic vocalizations. PLoS ONE, 10(8). 10.1371/journal.pone.0133251

Gaub, S., Fisher, S. E., & Ehret, G. (2016). Ultrasonic vocalizations of adult male Foxp2-mutant mice: Behavioral contexts of arousal and emotion. *Genes*, Brain and Behavior, 15(2). 10.1111/gbb.12274

Ghasemahmad, Z., Mrvelj, A., Panditi, R., Sharma, B., Perumal, D., & Wenstrup, J. J. (n.d.). Emotional Vocalizations Alter Behaviors and 2 Neurochemical Release into the Amygdala. 10.1101/2022.07.02.498564

Gill, S. A., & Bierema, A. M. K. (2013). On the Meaning of Alarm Calls: A Review of Functional Reference in Avian Alarm Calling. In Ethology (Vol. 119, Issue 6). 10.1111/eth.12097

Gourbal, B. E. F., Barthelemy, M., Petit, G., & Gabrion, C. (2004). Spectrographic analysis of the ultrasonic vocalisations of adult male and female BALB/c mice. Naturwissenschaften, 91(8). 10.1007/s00114-004-0543-7

Grimsley, J. M. S., Hazlett, E. G., & Wenstrup, J. J. (2013). Coding the meaning of sounds: Contextual modulation of auditory responses in the basolateral amygdala. Journal of Neuroscience, 33(44). 10.1523/JNEUROSCI.2205-13.2013

Grimsley, J. M. S., Monaghan, J. J. M., & Wenstrup, J. J. (2011). Development of social vocalizations in mice. PLoS ONE, 6(3). 10.1371/journal.pone.0017460

Grimsley, J. M. S., Sheth, S., Vallabh, N., Grimsley, C. A., Bhattal, J., Latsko, M., Jasnow, A., & Wenstrup, J. J. (2016). Contextual Modulation of Vocal Behavior in Mouse: Newly Identified 12 kHz “Mid-Frequency” Vocalization Emitted during Restraint. Frontiers in Behavioral Neuroscience, 10(March), 1–14. 10.3389/fnbeh.2016.00038

Guo, W., Chambers, A. R., Darrow, K. N., Hancock, K. E., Shinn-Cunningham, B. G., & Polley, D. B. (2012). Robustness of cortical topography across fields, laminae, anesthetic states, and neurophysiological signal types. Journal of Neuroscience, 32(27). 10.1523/JNEUROSCI.0065-12.2012

Hanson, J. L., & Hurley, L. M. (2012). Female presence and estrous state influence mouse ultrasonic courtship vocalizations. PLoS ONE, 7(7). 10.1371/journal.pone.0040782

Holmstrom, L., Roberts, P. D., & Portfors, C. v. (2007). Responses to social vocalizations in the inferior colliculus of the mustached bat are influenced by secondary tuning curves. Journal of Neurophysiology, 98(6). 10.1152/jn.00638.2007

Holy, T. E., & Guo, Z. (2005). Ultrasonic songs of male mice. PLoS Biology, 3(12). 10.1371/journal.pbio.0030386

Hurley, L. M., & Pollak, G. D. (2005). Serotonin modulates responses to species-specific vocalizations in the inferior colliculus. *Journal of Comparative Physiology A: Neuroethology, Sensory*, Neural, and Behavioral Physiology, 191(6). 10.1007/s00359-005-0623-y

Issa, J. B., Haeffele, B. D., Agarwal, A., Bergles, D. E., Young, E. D., & Yue, D. T. (2014). Multiscale Optical Ca2+ Imaging of Tonal Organization in Mouse Auditory Cortex. Neuron, 83(4). 10.1016/j.neuron.2014.07.009

Ito, T., & Malmierca, M. S. (2018). Neurons, Connections, and Microcircuits of the Inferior Colliculus. 10.1007/978-3-319-71798-2_6

Keesom, S. M., & Hurley, L. M. (2016). Socially induced serotonergic fluctuations in the male auditory midbrain correlate with female behavior during courtship. Journal of Neurophysiology, 115(4). 10.1152/jn.00742.2015

Kim, H., & Bao, S. (2013). Experience-dependent overrepresentation of ultrasonic vocalization frequencies in the rat primary auditory cortex. Journal of Neurophysiology, 110(5). 10.1152/jn.00230.2013

Klug, A., Bauer, E. E., Hanson, J. T., Hurley, L., Meitzen, J., & Pollak, G. D. (2002). Response selectivity for species-specific calls in the inferior colliculus of Mexican free-tailed bats is generated by inhibition. Journal of Neurophysiology, 88(4). 10.1152/jn.2002.88.4.1941

Kössl, M., & Vater, M. (1985). The cochlear frequency map of the mustache bat, Pteronotus parnellii. Journal of Comparative Physiology A, 157(5). 10.1007/BF01351362

Leroy, S. A., & Wenstrup, J. J. (2000). Spectral integration in the inferior colliculus of the mustached bat. Journal of Neuroscience, 20(22). 10.1523/jneurosci.20-22-08533.2000

Liu, R. C., & Schreiner, C. E. (2007). Auditory cortical detection and discrimination correlates with communicative significance. PLoS Biology, 5(7). 10.1371/journal.pbio.0050173

Lyzwa, D., Herrmann, J. M., & Wörgötter, F. (2016). Natural vocalizations in the mammalian inferior colliculus are broadly encoded by a small number of independent multi-units. Frontiers in Neural Circuits, 9(FEB2016). 10.3389/fncir.2015.00091

Maggio, J. C., Maggio, J. H., & Whitney, G. (1983). Experience-based vocalization of male mice to female chemosignals. Physiology and Behavior, 31(3). 10.1016/0031-9384(83)90186-5

Maggio, J. C., & Whitney, G. (1985). Ultrasonic vocalizing by adult female mice (Mus musculus). Journal of Comparative Psychology (Washington, D.C. : 1983), 99(4). 10.1037/0735-7036.99.4.420

Malmierca, M. S., Izquierdo, M. A., Cristaudo, S., Hernández, O., Pérez-González, D., Covey, E., & Oliver, D. L. (2008). A discontinuous tonotopic organization in the inferior colliculus of the rat. Journal of Neuroscience, 28(18). 10.1523/JNEUROSCI.0238-08.2008

Matsumoto, Y. K., & Okanoya, K. (2016). Phase-specific vocalizations of male mice at the initial encounter during the courtship sequence. PLoS ONE, 11(2). 10.1371/journal.pone.0147102

Mayko, Z. M., Roberts, P. D., & Portfors, C. v. (2012). Inhibition shapes selectivity to vocalizations in the inferior colliculus of awake mice. Frontiers in Neural Circuits, OCTOBER 2012. 10.3389/fncir.2012.00073

McLean, A. C., Valenzuela, N., Fai, S., & Bennett, S. A. L. (2012). Performing vaginal lavage, crystal violet staining, and vaginal cytological evaluation for mouse estrous cycle staging identification. Journal of Visualized Experiments, 67. 10.3791/4389

Mellott, J. G., Foster, N. L., Ohl, A. P., & Schofield, B. R. (2014). Excitatory and inhibitory projections in parallel pathways from the inferior colliculus to the auditory thalamus. Frontiers in Neuroanatomy, 8(November), 1–11. 10.3389/fnana.2014.00124

Neunuebel, J. P., Taylor, A. L., Arthur, B. J., & Roian Egnor, S. E. (2015). Female mice ultrasonically interact with males during courtship displays. ELife, 4(MAY). 10.7554/eLife.06203

Niemczura, A. C., Grimsley, J. M., Kim, C., Alkhawaga, A., Poth, A., Carvalho, A., & Wenstrup, J. J. (2020). Physiological and Behavioral Responses to Vocalization Playback in Mice. Frontiers in Behavioral Neuroscience, 14. 10.3389/fnbeh.2020.00155

Ostwald, J. (1984). Tonotopical organization and pure tone response characteristics of single units in the auditory cortex of the Greater Horseshoe Bat. Journal of Comparative Physiology A, 155(6). 10.1007/BF00611599

Petersen, C. L., & Hurley, L. M. (2017). Putting it in context: Linking auditory processing with social behavior circuits in the vertebrate brain. Integrative and Comparative Biology, 57(4). 10.1093/icb/icx055

Polese, A. G., Nigam, S., & Hurley, L. M. (2021). 5-HT1A Receptors Alter Temporal Responses to Broadband Vocalizations in the Mouse Inferior Colliculus Through Response Suppression. Frontiers in Neural Circuits, 15. 10.3389/fncir.2021.718348

Portfors, C. V. (2007). Types and functions of ultrasonic vocalizations in laboratory rats and mice. In Journal of the American Association for Laboratory Animal Science (Vol. 46, Issue 1).

Portfors, C. V. (2018). Processing of Ultrasonic Vocalizations in the Auditory Midbrain of Mice. In Handbook of Behavioral Neuroscience (Vol. 25). 10.1016/B978-0-12-809600-0.00007-X

Portfors, C. V., & Felix, R. A. (2005). Spectral integration in the inferior colliculus of the CBA/CaJ mouse. Neuroscience, 136(4). 10.1016/j.neuroscience.2005.08.031

Portfors, C. V., & Perkel, D. J. (2014). The role of ultrasonic vocalizations in mouse communication. In Current Opinion in Neurobiology (Vol. 28). 10.1016/j.conb.2014.07.002

Portfors, C. V., & Roberts, P. D. (2014). Mismatch of structural and functional tonotopy for natural sounds in the auditory midbrain. Neuroscience, 258. 10.1016/j.neuroscience.2013.11.012

Portfors, C. V., Roberts, P. D., & Jonson, K. (2009). Over-representation of species-specific vocalizations in the awake mouse inferior colliculus. Neuroscience, 162(2). 10.1016/j.neuroscience.2009.04.056

Ronald, K. L., Zhang, X., Morrison, M. V., Miller, R., & Hurley, L. M. (2020). Male mice adjust courtship behavior in response to female multimodal signals. PLoS ONE, 15(4). 10.1371/journal.pone.0229302

Sangiamo, D. T., Warren, M. R., & Neunuebel, J. P. (2020). Ultrasonic signals associated with different types of social behavior of mice. Nature Neuroscience, 23(3). 10.1038/s41593-020-0584-z

Scattoni, M. L., Crawley, J., & Ricceri, L. (2009). Ultrasonic vocalizations: A tool for behavioural phenotyping of mouse models of neurodevelopmental disorders. In Neuroscience and Biobehavioral Reviews (Vol. 33, Issue 4). 10.1016/j.neubiorev.2008.08.003

Schofield, B. R., & Hurley, L. (2018). Circuits for Modulation of Auditory Function. In D. Oliver, N. Cant, R. Fay, & A. Popper (Eds.), The Mammalian Auditory Pathways. Springer Handbook of Auditory Research (Vol. 65, pp. 235–265). Springer.

Schuller, G., & Pollak, G. (1979). Disproportionate frequency representation in the inferior colliculus of doppler-compensating Greater Horseshoe bats: Evidence for an acoustic fovea. Journal of Comparative Physiology □ A, 132(1). 10.1007/BF00617731

Sewell, G. D. S. (1972). Ultrasound and mating behaviour in rodents with some observations on other behavioural situations. Journal of Zoology, 168(2). 10.1111/j.1469-7998.1972.tb01345.x

Shepard, K. N., Lin, F. G., Zhao, C. L., Chong, K. K., & Liu, R. C. (2015). Behavioral relevance helps untangle natural vocal categories in a specific subset of core auditory cortical pyramidal neurons. Journal of Neuroscience, 35(6). 10.1523/JNEUROSCI.3803-14.2015

Stiebler, I., & Ehret, G. (1985). Inferior colliculus of the house mouse. I. A quantitative study of tonotopic organization, frequency representation, and tone-threshold distribution. Journal of Comparative Neurology, 238(1). 10.1002/cne.902380106

Stiebler, I., Neulist, R., Fichtel, I., & Ehret, G. (1997). The auditory cortex of the house mouse: Left-right differences, tonotopic organization and quantitative analysis of frequency representation. *Journal of Comparative Physiology - A Sensory*, Neural, and Behavioral Physiology, 181(6). 10.1007/s003590050140

Suga, N., & Jen, P. H. S. (1976). Disproportionate tonotopic representation for processing CF-FM sonar signals in the mustache bat auditory cortex. Science, 194(4264). 10.1126/science.973140

Šuta, D., Kvašňák, E., Popelář, J., & Syka, J. (2003). Representation of Species-Specific Vocalizations in the Inferior Colliculus of the Guinea Pig. Journal of Neurophysiology, 90(6). 10.1152/jn.01175.2002

Taberner, A. M., & Liberman, M. C. (2005). Response properties of single auditory nerve fibers in the mouse. Journal of Neurophysiology, 93(1). 10.1152/jn.00574.2004

Tsukano, H., Horie, M., Bo, T., Uchimura, A., Hishida, R., Kudoh, M., Takahashi, K., Takebayashi, H., & Shibuki, K. (2015). Delineation of a frequency-organized region isolated from the mouse primary auditory cortex. J Neurophysiol, 113, 2900–2920. 10.1152/jn.00932.2014.-The

Vater, M., Feng, A. S., & Betz, M. (1985). An HRP-study of the frequency-place map of the horseshoe bat cochlea: Morphological correlates of the sharp tuning to a narrow frequency band. Journal of Comparative Physiology A, 157(5). 10.1007/BF01351361

Wenstrup, J. J. (2005). The Tectothalamic System. In J. A. Winer & C. E. Schreiner (Eds.), The Inferior Colliculus (pp. 200–230). Springer.

Wenstrup, J. J., Nataraj, K., & Sanchez, J. T. (2012). Mechanisms of spectral and temporal integration in the mustached bat inferior colliculus. Frontiers in Neural Circuits, OCTOBER 2012. 10.3389/fncir.2012.00075

Williams, W. O., Riskin, D. K., & Mott, K. M. (2008). Ultrasonic sound as an indicator of acute pain in laboratory mice. Journal of the American Association for Laboratory Animal Science, 47(1).

Winer, J. A., & Schreiner, C. E. (2005). The inferior colliculus. In The Inferior Colliculus. 10.1007/b138578

Witzany, G. (2013). Biocommunication of animals. In Biocommunication of Animals. 10.1007/978-94-007-7414-8

Woolley, S. M. N., & Portfors, C. V. (2013). Conserved mechanisms of vocalization coding in mammalian and songbird auditory midbrain. In Hearing Research (Vol. 305, Issue 1). 10.1016/j.heares.2013.05.005

Zook, J. M., Winer, J. A., Pollak, G. D., & Bodenhamer, R. D. (1985). Topology of the central nucleus of the mustache bat’s inferior colliculus: Correlation of single unit properties and neuronal architecture. Journal of Comparative Neurology, 231(4). 10.1002/cne.902310410

